# A database of digital line drawings that depict expected and unexpected action-place relationships

**DOI:** 10.1101/2024.06.22.600189

**Authors:** Sabiha Tezcan Aydemir, Seray Tezcan Umul, Burcu A. Urgen

## Abstract

In the present study, we introduce an image database created with a simple digital line-drawing tool to represent the expected and unexpected action-place relationships. This database consists of 70 drawings. We validated the dataset with 207 participants. They evaluated the actions, places, and the probability of an action taking place in the respective location. The comprehensibility of each drawing was evaluated using three measures: H-statistics (entropy), which is a measure of the uncertainty of the definition; the definition similarity percentages which is a measure of the naming agreement that is consistent among participants; and the mean probability of the depicted action taking place in the depicted location. Each drawing includes different agents and environments, providing researchers with the opportunity to use this dataset in various fields of cognitive neuroscience, including visual recognition, memory, predictive coding, and novelty detection.

## Introduction

In psychology and neuroscience studies, various types of visual stimuli are used to evaluate behavioral and neural responses. These materials can include images, videos, drawings created by professionals or designed through artificial intelligence (AI), virtual reality, and other purpose-built visual materials that can be compiled from online sources (Parsons, 2015; Bucci Mansilla et al., 2021; Gobet et al., 2019). While creating these stimuli, not only the generation techniques but also the content and context of the materials are of great importance. For example, items consisting of some meaningless shapes provide less ecologically valid and limited data, while semantically meaningful, complex visual materials can generate more detailed behavioral and neural responses (Shaojie et al., 2022, Risko et al., 2012). Also, the elements in these tests can vary depending on the fundamental perceptual or cognitive function that is being assessed. While meaningless shapes can be sufficient when evaluating low-level visual functions (Barnett et. al, 2016; Quinlan et. al, 2012), more complex and semantically meaningful images such as objects, faces and scenes can be used in tasks involving visual recognition, memory, language, and associative learning (Barry et. al., 1997; Meyer, 2020; Düzel et. al, 2018; Chen et. al, 2021; Suet et al. 2012; Bunzeck and Duzel.2006; Kok et al. 2022; Hoef et al. 2024).

Face databases are at the forefront of the materials used as semantically meaningful items (Workman et al., 2021). Facial recognition has been evaluated in neuroscience and psychology for so long that quite extensive databases have been created in this area, and researchers can access different groups of items according to their target variables and research questions (Stan et al., 2005; Khushi et al., 2022). However, evaluations that are conducted with isolated images or drawings may be limited without having contextual information (Calder et al., 2005). This condition is relevant not only for isolated face databases but also for isolated scene and object databases as another group of semantically meaningful items, regardless of their size (Kriegeskorte et al., 2008; Baker et al., 2019; Torralba et al., 2010; Torralba et al., 2017; Zitnick et. al, 2014). On the other hand, incorporating a diverse range of semantically rich elements can lead to a better understanding of how the brain integrates various types of information to form coherent perceptions of the environment and can help researchers develop more comprehensive models of neural processing that are more applicable to real-world scenarios (Cicourel, 2005; Gordon et al., 2019). One such effort is the development of databases that include materials with object-scene relationships (Suet et al. 2012; Hoef et al.2024; Düzel et. al. 2018; Evans et al. 2022; Brandman et al. 2022). In these materials, researchers usually create congruent/expected and incongruent/unexpected situations according to statistical regularities of the environment and this provides researchers the opportunity to evaluate the detection of novelty, effects of surprise, predictive processing, language, and many other functions. For example, the Berlin Object Recognition Database is a rich database containing two groups of items, where the relationship between scene and object is represented. In this database, the photographing of the objects in expected and unexpected places provide much richer evaluation opportunities compared to basic facial, object, or scene images (Mohr et al., 2016).

While objects and their relationship with the environment have been extensively studied from a methodological point of view, there is limited research on actions and their relationship with the environment. Most of the action databases are created to lead the work in action observation studies in cognitive neuroscience (Lee Masson and Op de Beeck, 2018; Urgen et al., 2023; Georgiev et al 2023). While these databases provide a rich set of images or videos, they usually focus on a limited class of naturalistic actions (e.g. manipulation or social interaction) and fall short of depicting action-environment relationships in a systematic manner. Especially, given the increasing interest in predictive processing in cognitive neuroscience in recent years with a focus on actions (Kilner and Friston, 2007; Saygin et al., 2012; Urgen et al., 2018; Elmas et al., 2023, Uckan et al., 2024), there is a need for a database that includes visual items that depict congruent/expected and incongruent/unexpected action-environment relationships.

The aim of the present study is to create a database of actions that depict expected and unexpected relationships with the environment where the action takes place. To this end, the images are created in a controlled manner using digital line drawings, and H-statistic scores (entropy) are calculated to estimate the understandability of the images. We believe that this database can be used by researchers studying vision, memory, language, novelty detection, and predictive processing, and provide an important resource to examine how expectations influence perception and cognition.

## METHOD

### Participants

207 participants (142 female; Mean age = 36.40 years, *SD* =11.14) were included through online recruiting platform Qualtrics. The online Survey protocols in this study were approved by the Ethics Committee for Research with Human Participants of Bilkent University. All participants included in the study were over 18 years old, and they read and signed the online informed consent form before starting the surveys. Participants reported normal or corrected to-normal vision.

### Materials

There was a total of 72 visual elements, with 36 of them containing expected action-place relationships (e.g., a drawing of a person cooking in the kitchen), and 36 containing unexpected action-place relationships (e.g., a drawing of a person cooking in the bedroom). In this setup, for example, a drawing of a person cooking in the kitchen is considered to be an expected stimulus in terms of the relationship between the action and where it takes place and overlaps with the participant’s past world exposures, while a picture of someone cooking in the bedroom is thought to be perceived as a novel/unexpected stimulus. Before preparing these visual elements, a ’pre-study’ was conducted to identify the actions and place elements that would be drawn.

### Pre-study

For defining the actions and places to be drawn, a separate preliminary online survey was conducted to ensure that the items to be designed were the ones commonly exposed by the community. Qualtrics was used to compile open-ended questions regarding the actions to be drawn and the places in our daily lives, where these actions are expected or unexpected to perform. Following the survey’s distribution on social media platforms, data from 336 participants (206 female, Mean age=36.3 years, SD=7.2) was collected. Each participant needed 10 to 15 minutes to complete this survey.

According to the survey results, a total of 1562 action ideas were collected from all participants. These actions were then grouped into 90 action categories according to some criteria. For actions that would look similar when depicted, for instance, “walking” and “going for a walk” were grouped under the title “walking” to form a single action category, and “having breakfast,” “eating,” and “snacking” were included in the “eating” action category. However, actions that would look different when depicted, such as playing basketball or playing soccer, were grouped as separate actions, and a generic action group like ’doing sports’ was not created. In addition, some actions indicated by a single participant constituted the “other” category. Then, a total of 60 action categories were identified after excluding the actions that were difficult to express through drawing, such as crying, breathing, and helping each other. Among these 60 depictable action categories, we prefer the action-place relationships that are unexpected based on physical regularities rather than semantic regularities. This choice was made specifically to facilitate a better understanding when depicted and to create a more homogeneous test. For example, swimming on a carpet at home is a highly unexpected situation due to its physical difficulty. However, sleeping in the office is not as impossible as swimming on a carpet and is primarily socially unexpected. Considering these situations, of the 60 depictable ideas, 24 had to be eliminated and the remaining 36 ideas were utilized. Finally, it was decided to draw a total of 72 elements, including 36 expected-routine action-place relationships and 36 unexpected-new action-place relationships. All action categories and the drawn ones are shown in **Supplementary material.1** file.

### Design of the images

Once the actions and places were decided upon, one expected and one unexpected place was prepared for each action. One of the authors, who is a professional artist used the Procreate tool to create the black and white drawings as 3000*3000 pixels png files (Savage Interactive, 2024). The other two authors evaluated each item to determine whether the image was appropriate in terms of expected and unexpected action-place relationships. In these images, every actor and scene were unique for each item. The list of all items with their brief descriptions is provided in Table 1. Figure 1 shows four sample images from this list (two actions with expected and unexpected contexts).

**Figure 1.**
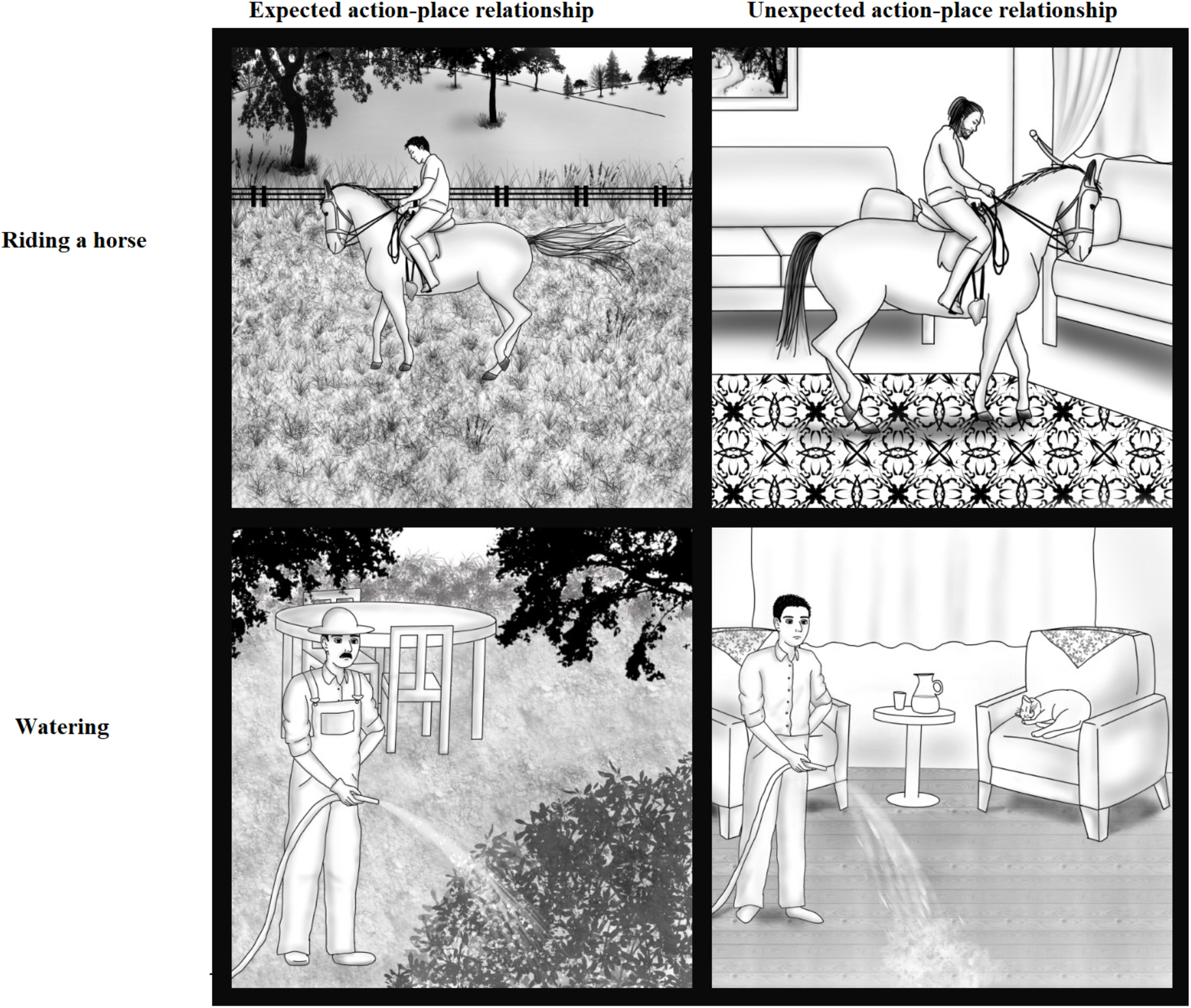
Sample drawings from the database that depict expected and unexpected action-place relationships.

**Table 1.** Brief descriptions of the images.

| No. | Action & Type | Description |
| --- | --- | --- |
| 1 | Knitting - expected | A woman is knitting on a sofa in the living room. |
| 2 | Driving - expected | A man is driving a car on a road. |
| 3 | Putting on makeup - expected | A woman is putting on makeup in a bedroom in front of a mirror. |
| 4 | Climbing - expected | A man is climbing a mountain. |
| 5 | Swimming - expected | A person is swimming in a pool. |
| 6 | Playing basketball - expected | A man is playing basketball on a basketball court. |
| 7 | Riding a horse - expected | A man is riding a horse in a field. |
| 8 | Vacuuming - expected | A woman is vacuuming in a living room. |
| 9 | Skating - expected | A woman is skating on a pavement. |
| 10 | Brushing teeth - expected | A woman is brushing her teeth in a bathroom in front of a mirror. |
| 11 | Picking flowers - expected | A woman is picking flowers in a meadow. |
| 12 | Working on a PC - expected | A man is working or studying on a PC in a study room. |
| 13 | Cutting grass - expected | A man is cutting grass in a garden. |
| 14 | Watering - expected | A man is watering plants in a garden. |
| 15 | Shaving - expected | A man is shaving in a bathroom in front of a mirror. |
| 16 | Skiing - expected | A person is skiing on a snowy mountain with ski equipment. |
| 17 | Jumping - expected | A woman is jumping on a trampoline in a garden. |
| 18 | Examining - expected | A doctor is examining a patient in a clinic. |
| 19 | Operating - expected | Two doctors are operating on someone in an operating room. |
| 20 | Ironing - expected | A woman is ironing in a living room. |
| 21 | Eating - expected | A woman is eating her meal at a table in the kitchen. |
| 22 | Fishing - expected | A man is fishing by the sea. |
| 23 | Withdrawing money - expected | A woman is withdrawing money from an ATM on a street. |
| 24 | Reading - expected | Two women are reading books in a library. |
| 25 | Walking the dog - expected | A woman is walking the dog on a road. |
| 26 | Sleeping - expected | A woman is sleeping in a bed in the bedroom. |
| 27 | Praying - expected | A man is praying in a mosque. |
| 28 | Walking - expected | A man is walking on a pavement. |
| 29 | Cooking - expected | A woman is cooking in a professional kitchen. |
| 30 | Jumping rope - expected | A boy is jumping rope in a park. |
| 31 | Starting a fire - expected | A man is starting a fire at a campsite. |
| 32 | Playing piano - expected | A man is playing a piano in a concert hall. |
| 33 | Getting a haircut - expected | A woman is getting a haircut in a hair salon. |
| 34 | Biking - expected | A woman is biking in a park. |
| 35 | Sunbathing - expected | A woman is sunbathing near a pool wearing swimsuits. |
| 36 | Shopping - expected | A man is shopping in a market. |
| 37 | Knitting - unexpected | A woman is knitting under the sea. |
| 38 | Driving - unexpected | A man is driving a car on the sea. |
| 39 | Putting on makeup - unexpected | A woman is putting on makeup on the floor of a gym. |
| 40 | Climbing - unexpected | A man is climbing a wardrobe in a bedroom with climbing equipment. |
| 41 | Swimming - unexpected | A man is swimming on a carpet in a living room wearing snorkel and swim goggles. |
| 42 | Playing basketball - unexpected | A man is playing basketball in a field near a tractor. |
| 43 | Riding a horse - unexpected | A man is riding a horse in a living room. |
| 44 | Vacuuming - unexpected | A woman is vacuuming leaves in a garden. |
| 45 | Skating - unexpected | A woman is skating on the sand at the seaside. |
| 46 | Brushing teeth - unexpected | A man is brushing his teeth in a subway. |
| 47 | Picking flowers - unexpected | A woman is picking flowers in a snowy area. |
| 48 | Working on a PC - unexpected | A man is working or studying on a PC in a Turkish bath. |
| 49 | Cutting grass - unexpected | A man is cutting grass in a large bedroom dressed as a gardener. |
| 50 | Watering - unexpected | A man is watering the floor in a living room. |
| 51 | Shaving - unexpected | A man is shaving on the road in his pajamas. |
| 52 | Skiing - unexpected | A person is skiing in a desert with ski equipment. |
| 53 | Jumping - unexpected | A woman is jumping on a moving bus. |
| 54 | Examining - unexpected | A doctor is examining a patient in front of a greengrocer. |
| 55 | Operating - unexpected | Two doctors are operating on someone in a repair shop. |
| 56 | Ironing - unexpected | A woman is ironing on a table in a restaurant. |
| 57 | Eating - unexpected | A woman is eating her meal in a bathtub in the bathroom. |
| 58 | Fishing - unexpected | A man is fishing in a bus terminal. |
| 59 | Withdrawing money - unexpected | A woman is withdrawing money from an ATM in a living room. |
| 60 | Reading - unexpected | A man is reading a book in a refrigerator. |
| 61 | Walking the dog - unexpected | A woman is walking the dog on the roof of a house. |
| 62 | Sleeping - unexpected | A woman is sleeping in a car on the street. |
| 63 | Praying - unexpected | A man is praying in a toilet. |
| 64 | Walking - unexpected | A man is walking on a table in the kitchen. |
| 65 | Cooking - unexpected | A woman is cooking in a bedroom. |
| 66 | Jumping rope - unexpected | A woman is jumping rope in a store. |
| 67 | Starting a fire - unexpected | A girl is starting a fire in an empty classroom. |
| 68 | Playing piano - unexpected | A man is playing a piano in the desert. |
| 69 | Getting a haircut - unexpected | A woman is getting a haircut in a butcher shop. |
| 70 | Biking - unexpected | A woman is biking in a living room. |
| 71 | Sunbathing - unexpected | A woman is sunbathing in a construction area wearing swimsuits. |
| 72 | Shopping - unexpected | A man is shopping in an emergency room of a hospital. |

## Main Study

### Procedure

Following the completion of the design of the images, an online survey was administered via Qualtrics to ensure that the actions, places, and action-place relationships depicted in the drawings were understood as intended. This survey was spread through similar social media platforms as the previous one, and data were collected from 207 adults (142 Female; Mean age: 36.40; SD: 11.14). The number of participants to be included in the survey was determined to be at least 100-150 participants, considering the participant numbers in similar studies (Brown et al., 1976; Souza et al., 2020). In the survey, three questions were asked for each image item. The first questions required the participants to name the actions and places in the images, respectively. In the final question, participants were asked to indicate the likelihood of encountering the depicted action in the drawn place using a slider ranging from 0 to 100. It took 30 to 45 minutes to complete the survey.

## Data Analysis

The statistical analysis and machine learning tool in MATLAB 2022b version was used to analyze and graph the data (MATLAB, 2022).

In the pre-study, the open-ended responses (five actions and five places for each participant) were assessed and categorized manually. Then, the mean and standard deviation values of the participants’ demographic data were calculated.

In the main study, the open-ended responses (the first two questions) of the survey were assessed and categorized manually. Subsequently, entropy analyses (H-statistic) were conducted on these categories, and the frequency of responses in the first category, referred to as the most frequently given answer, was calculated as a percentage. For the third question of the main study, probability percentage values were calculated as means and standard deviations.

### Response categorization of action and place names

Since the first two questions required open-ended answers, the responses needed to be qualitatively analyzed to determine whether the intended action and place were understood. In analyzing the responses both for actions and places, one or more groups were identified for each item: the first group always indicated the intended action or place category. For example, for the action of walking, responses like “walking", “strolling", and “taking a walk” were all included in the first group. The increasing group number typically represented a deviation from the intended response. For instance, in the image of a person jumping on a bus, naming the action as “jumping” was grouped in the first category, while related yet different actions such as “being happy”, or “dancing” formed the second category. On the other hand, the response “flying” was distinct from the other two groups and significantly different from the intended action, thus formed the third response category. The frequency of given responses across participants was also considered while creating these categories. For a response to form a separate category different from other ones, it was required that more than one participant provided that response. As mentioned previously, responses given by only one person have been included in the ’other’ category.

Another factor considered in separating the action response categories was how specifically the image depicted the action. For example, playing basketball requires quite specific equipment. Therefore, responses like playing with a ball or just playing, which are correct but less specific, were grouped in a separate category. This approach was followed because it conflicted with the initial manipulation done in the images. By initial manipulation, we mean the deliberate creation of expected or unexpected action-place relationships in the images. Accordingly, for instance, playing basketball in a farm field, which represents an unexpected action-space relationship, is practically impossible due to the lack of necessary equipment in the field. However, playing with a ball or just playing in the field are plausible actions. This is why answers mentioning “playing basketball” and “playing” have been placed into distinct categories.

The initial manipulation used in the drawing of the images has also been considered in the categorization of the place names in the images. Consequently, for example, the response “kitchen” to an image of someone reading a book inside a refrigerator (depicted as reading in a refrigerator in the kitchen) was not entirely incorrect, but it was distinguished from the response “inside the refrigerator.” The manipulation considered here makes reading inside the refrigerator an unexpected element. Thus, describing an image considered to contain an unexpected element such as reading a book in the kitchen actually indicates a situation that is not unexpected. In such cases, “kitchen” and “refrigerator” definitions have also formed different categories. In addition to this, in naming places, it was expected to specifically name places that contain quite specific objects. For example, for an image of a bathroom, only “bathroom” or “home” (if not contradictory to the initial manipulation) was included in the first category, while responses like “living room” or “kitchen” for this picture were separated. However, places like living room, dining room, and lounge, which have similar objects, were generally included in the same category.

### H-statistics and Name Agreement for action and place names

After the grouping of the responses to the open-ended questions, we calculated H-statistic for each image. The H-statistic, also known as entropy, is used to measure the level of uncertainty in identifying an image. This metric, based on a methodology from Snodgrass and Vanderwart (1980) and further developed by Rossion and Pourtois (2004), calculates entropy by considering the number of valid responses per image (excluding any invalid responses) and the proportion of participants who gave each response. As Lachman (1973) noted, the H-statistic quantifies the uncertainty in labeling an object. Basically, a low H-statistic indicates a high level of agreement among participants on the name of an image (a value of 0 means everyone agreed on the same name), while a high H-statistic indicates a low level of agreement, reflecting a wide variety of names given to the same image.

### Probability analysis

For the third question of the main study which asked about the probability of an action being taken in a respective place, we calculated the mean and standard deviation of the percentages across all participants for each image.

## RESULTS

The analysis of the third question in the main study revealed that most of the expected items had a value of 90% and above, while the unexpected items had a degree of 10% and below. However, the probability values for 9 items were between these values. In order to create a more homogenous set, these 9 items were redesigned, and the same survey was repeated only with these 9 items with 192 participants (140 women). We then merged the results of the main study with this final re-design study.

Figure 2 shows the mean probabilities (in terms of the action-place relationship) given for each image across all participants. The numbering of the images in this graph corresponds to Table 1. The images that contained unexpected action-place relationships generally had a probability average of 10% and below (mean of means=6.33), while those that were expected had an average above 90% (mean of means=94.82). However, despite further adjustments of the nine images, it was observed that a visual element containing one expected (“skating”) and one unexpected ("putting on make-up”) action-place relationship had a probability mean between these extreme values (See Figure 2, items marked with red). Consequently, these two images were excluded from the final set. As a result, we decided to use 70 elements, half expected and half unexpected.

**Figure 2.**
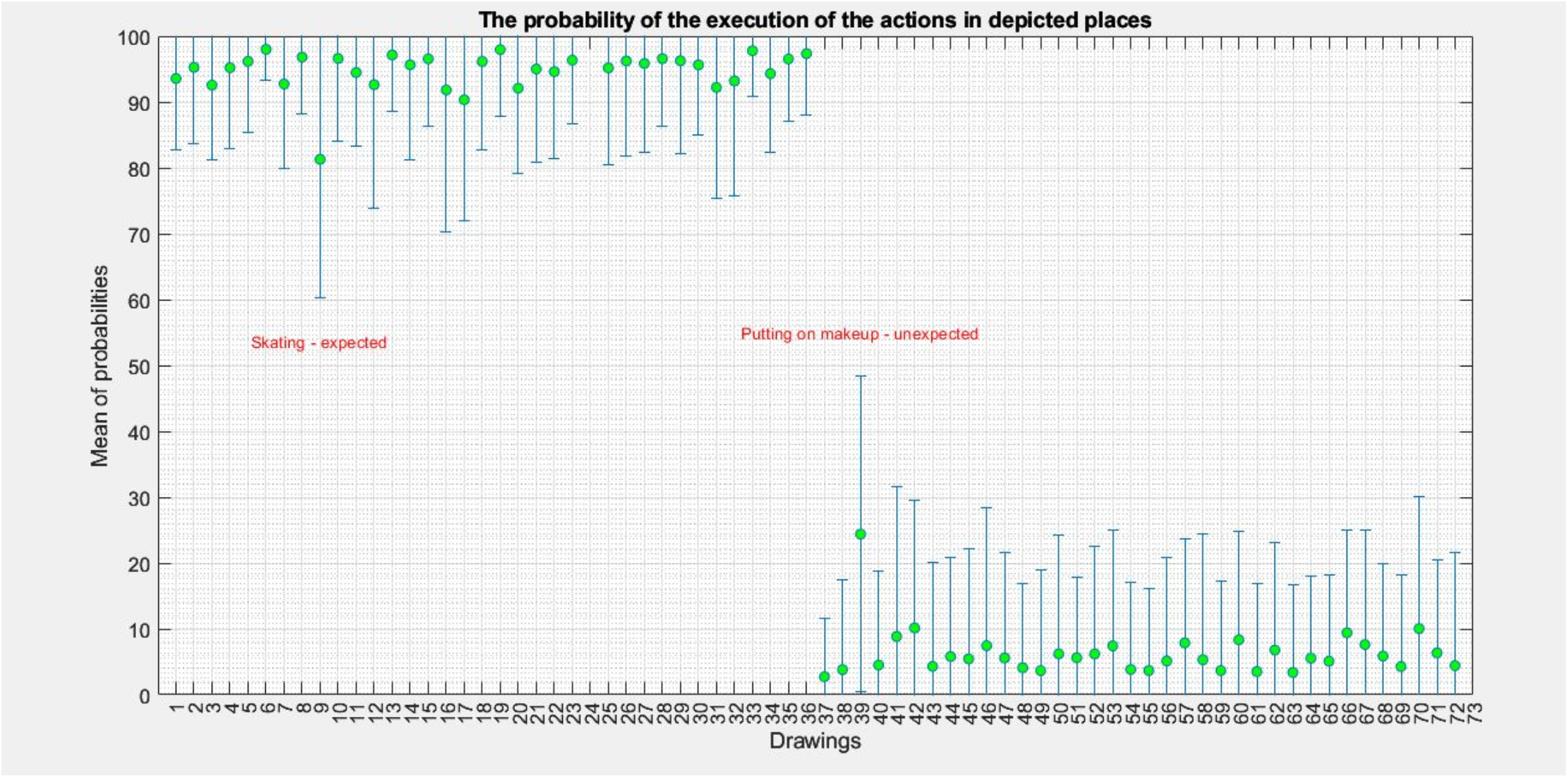
The probabilities of actions in the drawings taking place in the depicted location.

After categorizing the responses to the open-ended questions, we calculated the H-statistics for the expected and unexpected items. H-statistics for the action and place names together with the probability values of the action-place relationships are summarized in **Table 2 and Table 3**, **respectively.** The categories for the action and place names that contributed to the H-statistics were provided in **Supplementary Material 2.**

**Table 2.** Statistical Analysis and Name Agreement Results for Expected Images.

| Expected items | Action definition similarity | Action H-statistics | Place definition similarity | Place H-statistics | The probability of the actions takes place in that place (mean ±SD) |
| --- | --- | --- | --- | --- | --- |
| Knitting | 98.55% | 0.109 | 98.55 | 0.122 | 95.57±10.79 |
| Driving | 98.06% | 0.153 | 93.23 | 0.420 | 95.26±11.52 |
| Putting on makeup | 100.00% | 0 | 100.00% | 0 | 92.57±11.41 |
| Climbing | 99.50% | 0.044 | 99.03% | 0.078 | 95.20±12.35 |
| Swimming | 99.50% | 0.044 | 99.03% | 0.078 | 96.15±10.78 |
| Playing Basketball | 99.50% | 0.044 | 100.00% | 0 | 97.99±4.70 |
| Riding a horse | 100.00% | 0 | 99.03% | 0.078 | 92.73±7.62 |
| Vacuuming | 100.00% | 0 | 99.50% | 0.044 | 96.81±8.68 |
| Skating | 98.95% | 0.083 | 100% | 0 | 81.01±22.22 |
| Brushing teeth | 100.00% | 0 | 100.00% | 0 | 96.60±12.51 |
| Picking flowers | 100.00% | 0 | 99.50% | 0.044 | 94.47±11.18 |
| Starting a fire | 87.43% | 0.574 | 100.00% | 0 | 92.22±16.85 |
| Working on PC | 99.50% | 0.044 | 99.50% | 0.044 | 92.63±18.68 |
| Cutting the grasses | 100.00% | 0 | 100.00% | 0 | 97.14±8.53 |
| Watering | 100.00% | 0 | 100.00% | 0 | 95.64±14.35 |
| Shaving | 100.00% | 0 | 100.00% | 0 | 96.55±10.23 |
| Skiing | 99.50% | 0.044 | 97.58% | 0.164 | 91.82±21.49 |
| Jumping | 98.55% | 0.109 | 49.27% | 1.094 | 90.33±18.34 |
| Examining | 100.00% | 0 | 99.50% | 0.044 | 96.15±13.37 |
| Operating | 100.00% | 0 | 99.50% | 0.044 | 97.95±10.21 |
| Ironing | 100.00% | 0 | 98.03% | 0.078 | 92.09±13.02 |
| Eating | 100.00% | 0 | 100.00% | 0 | 95.00±14.18 |
| Fishing | 100.00% | 0 | 99.50% | 0.044 | 94.61±13.16 |
| Withdrawing money | 100.00% | 0 | 99.03% | 0.078 | 96.33±9.64 |
| Reading | 92.27% | 0.392 | 100.00% | 0 | 95.95±13.03 |
| Walking the dog | 96.13% | 0.236 | 100.00% | 0 | 95.18±14.61 |
| Sleeping | 100.00% | 0 | 100.00% | 0 | 96.23±14.41 |
| Praying | 100.00% | 0 | 100.00% | 0 | 95.85±13.49 |
| Walking | 100.00% | 0 | 99.50% | 0.044 | 96.57±10.19 |
| Cooking | 100.00% | 0 | 100.00% | 0 | 96.25±14.12 |
| Jumping rope | 99.50% | 0.044 | 100.00% | 0 | 95.63±10.69 |
| Playing a piano | 100.00% | 0 | 97.58% | 0.164 | 93.19±17.44 |
| Getting a haircut | 100.00% | 0 | 100.00% | 0 | 97.77±6.93 |
| Riding a bike | 100.00% | 0 | 100.00% | 0 | 94.30±12.01 |
| Sunbathig | 99.03% | 0.078 | 97.58% | 0.181 | 96.52±9.41 |
| Shopping | 98.06 | 0.137 | %100.00 | 0 | 97.34±9.30 |

**Table 3.** Statistical Analysis and Name Agreement Results for Unexpected Images.

| Unexpected items | Action definition similarity | Action H-statistics | Place definition similarity | Place H-statistics | The probability of the actions takes place in that place (mean ±SD) |
| --- | --- | --- | --- | --- | --- |
| Playing a piano | 100.00% | 0 | 98.43% | 0.130 | 5.85±14.65 |
| Withdrawing money | 99.50% | 0.044 | 96.60% | 0.213 | 3.67±13.60 |
| Starting a fire | 98.50% | 0.109 | 100.00% | 0 | 7.62±17.34 |
| Putting on makeup | 100.00% | 0 | 100.00% | 0 | 24.24± 24.98 |
| Cutting the grasses | 96.13% | 0.272 | 83.09% | 0.655 | 3.66±15.31 |
| Jumping rope | 100.00% | 0 | 98.06% | 0.137 | 9.37±16.18 |
| Sleeping | 100.00% | 0 | 89.37% | 0.488 | 6.79±16.36 |
| Walking the dog | 96.13% | 0.236 | 100.00% | 0 | 3.51±13.35 |
| Riding a horse | 100.00% | 0 | 100.00% | 0 | 4.31±15.83 |
| Knitting | 97.38% | 0.200 | 98.95% | 0.0839 | 2.73±9.15 |
| Ironing | 99.50% | 0.044 | 97.50% | 0.181 | 5.13±15.72 |
| Driving | 98.06 | 0.137 | 99.50% | 0.044 | 3.82±13.63 |
| Playing basketball | 97.50% | 0.164 | 99.50% | 0.044 | 9.93±19.39 |
| Climbing | 95.60% | 0.297 | 99.50% | 0.044 | 4.51±14.26 |
| Skiing | 100.00% | 0 | 98.50% | 0.109 | 6.19±16.25 |
| Operating | 99.03% | 0.088 | 100.00% | 0 | 3.67±12.48 |
| Eating | 99.03% | 0.078 | 100.00% | 0 | 7.87±15.71 |
| Picking flowers | 98.50% | 0.109 | 98.06% | 0.137 | 5.58±15.93 |
| Skating | 95.16% | 0.344 | 99.03% | 0.078 | 5.44±16.73 |
| Jumping | 95.16% | 0.321 | 85.02% | 0.722 | 7.42±17.52 |
| Shaving | 98.50% | 0.109 | 100.00% | 0 | 5.61±12.15 |
| Working on PC | 100.00% | 0 | 100.00% | 0 | 4.09±12.78 |
| Brushing teeth | 99.03% | 0.078 | 100.00% | 0 | 7.45±20.98 |
| Riding bike | 98.55% | 0.122 | 100.00% | 0 | 10.04±20.11 |
| Fishing | 98.50% | 0.109 | 99.03% | 0.078 | 5.32±19.16 |
| Vacuuming | 99.03% | 0.078 | 99.03% | 0.078 | 5.80±15.05 |
| Shopping | 96.13% | 0.277 | 100.00% | 0 | 4.43±17.26 |
| Cooking | 99.03% | 0.078 | 99.03% | 0.078 | 5.09±13.07 |
| Praying | 100.00% | 0 | 99.50% | 0.044 | 3.38±13.32 |
| Examining | 100.00% | 0 | 100.00% | 0 | 3.84±13.32 |
| Swimming | 71.00% | 1.329 | 99.03% | 0.088 | 8.85±15.12 |
| Getting a haircut | 100% | 0 | 97.38% | 0.193 | 4.28±14.58 |
| Walking | 98.42% | 0.131 | 91.62% | 0.415 | 5.53±12.91 |
| Reading | 98.95% | 0.083 | 92.14% | 0.396 | 8.31±17.21 |
| Sunbathing | 98.42% | 0.116 | 92.14% | 0.496 | 6.32±14.79 |
| Watering | 100.00% | 0 | 100.00% | 0 | 6.20±17.94 |

## Discussion

In this study, 72 simple digital line drawings were created, with half representing expected and the other half representing unexpected action-place relationships. While creating these images, the pre-study focused on actions and places commonly encountered in daily life. Different individuals and various locations were depicted in each drawing. After the design phase, the drawings were assessed by a large number of participants for validation. We then computed definition similarity percentages, which is a measure of naming agreement, H-statistics (entropy), which is a measure of uncertainty of the definitions for action and place names, and the mean probability values for the action-place relationships.

The results show that all action and place definitions have low H-statistics and high similarity percentages indicating agreement across individuals. Among all drawings, only one item had an H statistic value exceeding 1. Most of the similarity values exceed 95 percent, except for three specific actions and seven place definitions. When these exceptional items are examined further, similarity values below 95% in naming were observed only for very surprising, unexpected items (jumping on the bus, swimming in the living room) or for existing specific objects and locations that might not be known by everyone (such as a trampoline). This situation could be considered an indicator that the idea of dynamic actions can be represented with static elements, independent of the manipulation performed. To achieve this, even if the representation of the place where the action takes place is unexpected, the specific objects related to the action and the positions of the agents may have clarified the concept for participants.

Similar to the labeling of actions, the naming of places has also been facilitated by the presence and arrangement of certain objects, especially in expected images. In these expected images, only one place has a naming similarity percentage below 95%. Additionally, the presence of actions and agents in the pictures may have made it easier to recognize some places. For example, the presence of a person jumping on a trampoline or someone skiing on a snowy mountain can facilitate the definitions of the places. Without the jumping person, it might have been difficult to name a trampoline standing in a garden in a black-and-white image. The same could be valid for naming a snowy mountain in a black-and-white picture without seeing a person equipped with ski attire. In contrast, the situation was somewhat different for items involving unexpected action-place relationships. Six place items had naming percentages below 95%. Upon examining this naming diversity, it can be suggested that this variety exists because participants might have tried to make sense of the situation in the picture within a logical framework. For instance, some participants tried to rationalize the appearance of a butcher, which is an unexpected place for a haircut, by naming the place as a shopping mall with multiple stores where a butcher could be seen in the background or the garden of a hair salon.

These results indicate that the concept of dynamic action and its relationship with the place could be appropriately represented through static visual elements in this set of images. Even in elements containing unexpected action-location relationships, there were notably high levels of naming consistency, and the probability values distinctly diverged in a manner consistent with the manipulations performed. It is important to recognize that creating a semantically meaningful dataset can have some sensitive aspects. For instance, compiling a dataset from various internet sources rather than creating it according to the participants’ goals can pose significant challenges for neuroscience and psychology experiments. During the process of collecting visuals from various sources, procedures like image compression methods and graphic editing can lead to distortion and a decline in quality of the elements. This can significantly impact the uniformity of the dataset and result in deviations from typical viewing behavior (Wichmann, 2010). Given this situation, within the framework of this study, the dataset created entirely by a single professional using a single tool with basic line drawings is significantly advantageous in terms of homogeneity. It is evident that creating such datasets by a single individual is essential, not only for maintaining technical consistency but also for ensuring precise semantic manipulation. For instance, in existing research, a study assessed participants’ emotions and moral judgments by exposing them to images of faces that included anomalies, serving as unexpected yet semantically significant elements. These anomalies included dermatitis, injury scars, and congenital anomalies. The study discussed the challenges of interpreting evaluations made with these anomalous faces, which represented very different ideas and were not grouped within certain limits (Workman et al., 2021). From this perspective, it can be said that starting with a specific concept provided a significant advantage in the creation of our elements. In this way, repetition of certain ideas has been avoided, and the degrees of expectancy and unexpectedness have been created at similar levels, with few exceptions.

This database which is designed with precise semantic ideas can be useful in novelty detection studies. Because, in studies of novelty detection, an element containing a routine action-place relationship can be found familiar according to our past world knowledge, while an element containing an unexpected action-place relationship can be considered a novel item. This makes these novel elements particularly appropriate in assessing the detection of semantic novelty, which is rarely evaluated in novelty detection studies (Suet et al. 2012; Nianzu et al. 2021; Nentwitch et al. 2023). In previous novelty detection literature, studies on contextual novelty and associative novelty, which are typically conducted through repeated presentations of items or changes in item relationships, are more common (Daffner et al. 2001; Bastin et al. 2019; Miller et al. 2008; O’Brien et al. 2010; Putcha et al. 2011; Dickerson et al. 2005). This database allows for the creation of the concept of novelty independently of a specific test sequence, focusing on individual items.

Additionally, these elements, which include two groups of action-place relationships, can also be utilized in predictive coding studies. In the literature on predictive coding, the relationship between cues and stimuli often is constructed with the help of short-term memory is frequently used (Van Kesteren et al. 2012; Gaeta vd.1999; Pekkonen et al. 2001; Sanders vd. 2022). However, two groups of elements defined as expected and unexpected according to cues from our semantic memory knowledge can provide different insights in these assessments. This feature also makes the current test items more effective in vulnerable populations such as elderly individuals with dominant short-term memory deficits and people with Alzheimer’s disease dementia.

Visual elements that contain expected and unexpected relationships find use not only in neuroscience studies but are also extensively used in applications such as the AI-mediated grouping of visual elements. For example, elements of the Berlin Object Recognition Test, which include previously mentioned expected and unexpected object and location relationships, have also been used for this purpose (Mohr et al., 2016). The elements of this study are also used by researchers for similar purposes.

## Limitations

In this study, the total number of 70 elements created might be considered somewhat small compared to standard databases. However, during the phase of representing the concept of action through drawings, the need to ensure clarity led us to exclude many actions that are frequently performed in everyday life, which is the reason for this limitation. Additionally, considering the demographic characteristics of the survey participants who evaluated the images, the number of participants over the age of 50 was quite low, and all participants had at least a high school education. Therefore, careful interpretation of the results is necessary when using these test elements with participants who have a lower educational level and older participants.

In addition to all these, the widespread use of social media and the frequent use of surprising, unexpected scenes in the advertising industry can change the impact of elements representing the relationship between action and place that we describe as unexpected over time. For example, according to survey results, the image of ’putting on makeup at the gym,’ which was excluded from the research and prelabeled as an unexpected element, was not found as unexpected by the participants as other images. It was thought that this might be due to the impact created by social media content. Furthermore, the alignment of elements with expectations can vary across different societies. For instance, the image of ’skating on the sidewalk,’ which we thought could be an expected element, was also not found as expected by the participants and was excluded from the dataset. This situation was associated with the fact that skating is not common in the society where the research was conducted. In summary, it would be beneficial to consider this situation when using the dataset in different societies.

## CONCLUSIONS

Semantically meaningful visual elements can provide valuable data for psychology and neuroscience studies (Suet et al. 2012; Bunzeck and Duzel.2006; Düzel et. al, 2018; Kok et al. 2022; Hoef et al. 2024). Associations in these visual elements, such as object-location and person-object relationships, not only allow for various manipulations but also enrich the collected data. Utilizing the action-place relationship as one of these dual relationships can be a highly useful method when interpreting semantic information. In this context, we believe that the visual test, which includes manipulations of two different groups (expected and unexpected) and is created through a standard tool, could be an important test for future use in areas such as novelty detection, memory, language, and predictive coding in cognitive neuroscience.

## Declerations

**Funding:** No funding was received to assist with the preparation of this manuscript.

**Conflicts of interest/Competing interests:** Not applicable

**Ethics approval:** The questionnaire and methodology for this study was approved by the Ethics Committee of Bilkent University (16.03.2023/405) and was conducted in accordance with the ethical standards laid down in the 1964 Declaration of Helsinki and its later amendments or comparable ethical standards.

**Consent to participate:** Before participating in the online survey, participants read the study summary and indicated their understanding and willingness to participate by selecting a checkbox. They then proceeded to answer the survey questions.

**Consent for publication:** Informed consent was obtained from all individual participants included in the study.

**Availability of data and materials**: Stimuli and validation data will be made available on Open Science Framework upon publication of the paper.

**Code availability:** Not applicaple

## Authors’ contributions

Conceptualization: Sabiha Tezcan Aydemir, Seray Tezcan Umul, Burcu Ayşen Ürgen Methodology: Sabiha Tezcan Aydemir, Seray Tezcan Umul, Burcu Ayşen Ürgen Data Collection: Sabiha Tezcan Aydemir

Data Analysis: Sabiha Tezcan Aydemir

Writing - Original Draft: Sabiha Tezcan Aydemir

Writing - Review & Editing: Sabiha Tezcan Aydemir, Burcu Ayşen Ürgen Supervision: Burcu Ayşen Ürgen

## Acknowledgments and Funding Information

We would like to thank Prof. Dr. Erguvan Tuğba Özel Kızıl, Tuğçe Nur Pekçetin, and Gaye Aşkın for their valuable input on the study’s methodology and Prof. Hüseyin Boyacı for his great support. This study did not receive any external funding.

---

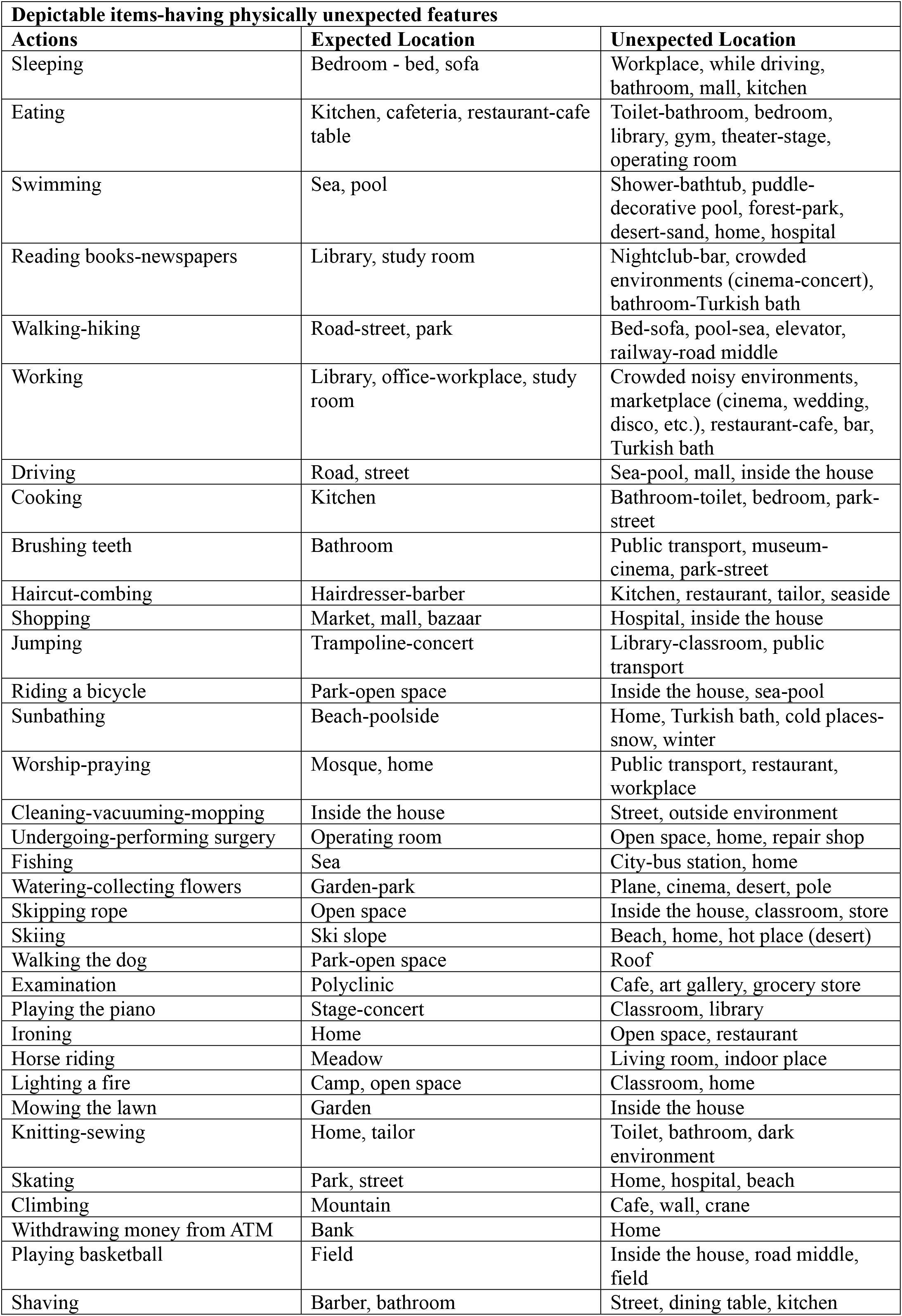
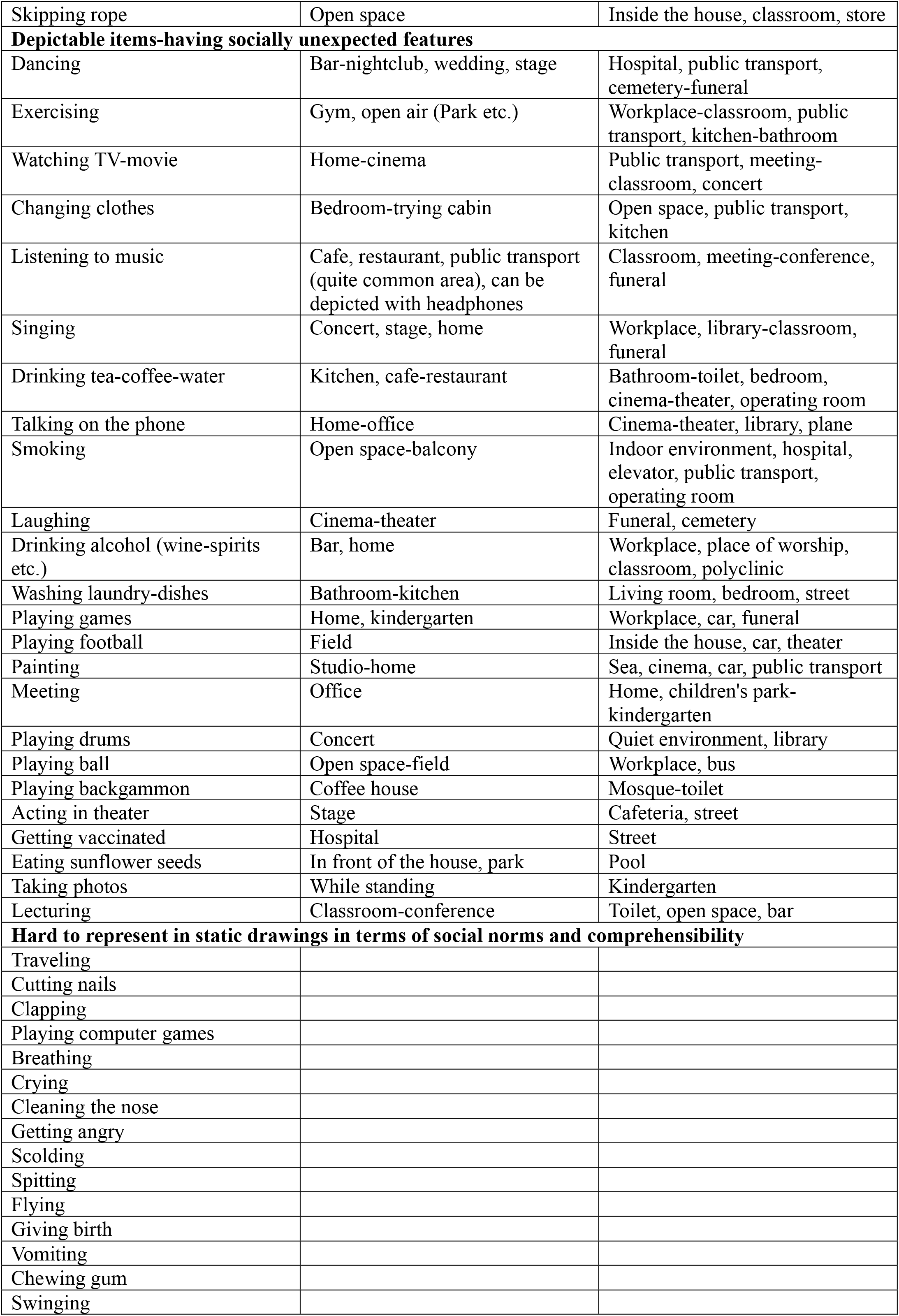
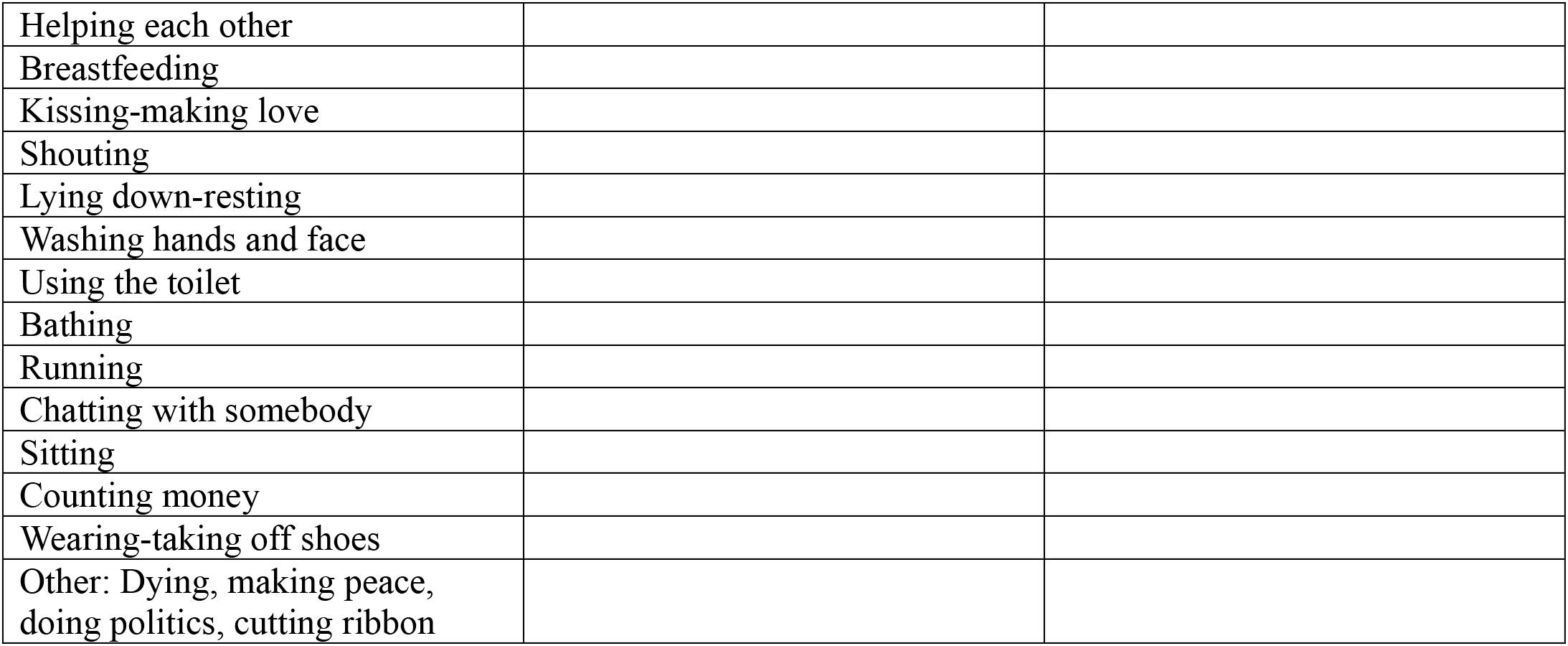

| **Depictable items-having physically unexpected features** |  |  |
| --- | --- | --- |
| **Actions** | **Expected Location** | **Unexpected Location** |
| Sleeping | Bedroom - bed, sofa | Workplace, while driving, bathroom, mall, kitchen |
| Eating | Kitchen, cafeteria, restaurant-cafe table | Toilet-bathroom, bedroom, library, gym, theater-stage, operating room |
| Swimming | Sea, pool | Shower-bathtub, puddle-decorative pool, forest-park, desert-sand, home, hospital |
| Reading books-newspapers | Library, study room | Nightclub-bar, crowded environments (cinema-concert), bathroom-Turkish bath |
| Walking-hiking | Road-street, park | Bed-sofa, pool-sea, elevator, railway-road middle |
| Working | Library, office-workplace, study room | Crowded noisy environments, marketplace (cinema, wedding, disco, etc.), restaurant-cafe, bar, Turkish bath |
| Driving | Road, street | Sea-pool, mall, inside the house |
| Cooking | Kitchen | Bathroom-toilet, bedroom, park-street |
| Brushing teeth | Bathroom | Public transport, museum-cinema, park-street |
| Haircut-combing | Hairdresser-barber | Kitchen, restaurant, tailor, seaside |
| Shopping | Market, mall, bazaar | Hospital, inside the house |
| Jumping | Trampoline-concert | Library-classroom, public transport |
| Riding a bicycle | Park-open space | Inside the house, sea-pool |
| Sunbathing | Beach-poolside | Home, Turkish bath, cold places-snow, winter |
| Worship-praying | Mosque, home | Public transport, restaurant, workplace |
| Cleaning-vacuuming-mopping | Inside the house | Street, outside environment |
| Undergoing-performing surgery | Operating room | Open space, home, repair shop |
| Fishing | Sea | City-bus station, home |
| Watering-collecting flowers | Garden-park | Plane, cinema, desert, pole |
| Skipping rope | Open space | Inside the house, classroom, store |
| Skiing | Ski slope | Beach, home, hot place (desert) |
| Walking the dog | Park-open space | Roof |
| Examination | Polyclinic | Cafe, art gallery, grocery store |
| Playing the piano | Stage-concert | Classroom, library |
| Ironing | Home | Open space, restaurant |
| Horse riding | Meadow | Living room, indoor place |
| Lighting a fire | Camp, open space | Classroom, home |
| Mowing the lawn | Garden | Inside the house |
| Knitting-sewing | Home, tailor | Toilet, bathroom, dark environment |
| Skating | Park, street | Home, hospital, beach |
| Climbing | Mountain | Cafe, wall, crane |
| Withdrawing money from ATM | Bank | Home |
| Playing basketball | Field | Inside the house, road middle, field |
| Shaving | Barber, bathroom | Street, dining table, kitchen |
| Skipping rope | Open space | Inside the house, classroom, store |
| **Depictable items-having socially unexpected features** |  |  |
| Dancing | Bar-nightclub, wedding, stage | Hospital, public transport, cemetery-funeral |
| Exercising | Gym, open air (Park etc.) | Workplace-classroom, public transport, kitchen-bathroom |
| Watching TV-movie | Home-cinema | Public transport, meeting-classroom, concert |
| Changing clothes | Bedroom-trying cabin | Open space, public transport, kitchen |
| Listening to music | Cafe, restaurant, public transport (quite common area), can be depicted with headphones | Classroom, meeting-conference, funeral |
| Singing | Concert, stage, home | Workplace, library-classroom, funeral |
| Drinking tea-coffee-water | Kitchen, cafe-restaurant | Bathroom-toilet, bedroom, cinema-theater, operating room |
| Talking on the phone | Home-office | Cinema-theater, library, plane |
| Smoking | Open space-balcony | Indoor environment, hospital, elevator, public transport, operating room |
| Laughing | Cinema-theater | Funeral, cemetery |
| Drinking alcohol (wine-spirits etc.) | Bar, home | Workplace, place of worship, classroom, polyclinic |
| Washing laundry-dishes | Bathroom-kitchen | Living room, bedroom, street |
| Playing games | Home, kindergarten | Workplace, car, funeral |
| Playing football | Field | Inside the house, car, theater |
| Painting | Studio-home | Sea, cinema, car, public transport |
| Meeting | Office | Home, children's park-kindergarten |
| Playing drums | Concert | Quiet environment, library |
| Playing ball | Open space-field | Workplace, bus |
| Playing backgammon | Coffee house | Mosque-toilet |
| Acting in theater | Stage | Cafeteria, street |
| Getting vaccinated | Hospital | Street |
| Eating sunflower seeds | In front of the house, park | Pool |
| Taking photos | While standing | Kindergarten |
| Lecturing | Classroom-conference | Toilet, open space, bar |
| **Hard to represent in static drawings in terms of social norms and comprehensibility** |  |  |
| Traveling |  |  |
| Cutting nails |  |  |
| Clapping |  |  |
| Playing computer games |  |  |
| Breathing |  |  |
| Crying |  |  |
| Cleaning the nose |  |  |
| Getting angry |  |  |
| Scolding |  |  |
| Spitting |  |  |
| Flying |  |  |
| Giving birth |  |  |
| Vomiting |  |  |
| Chewing gum |  |  |
| Swinging |  |  |
| Helping each other |  |  |
| Breastfeeding |  |  |
| Kissing-making love |  |  |
| Shouting |  |  |
| Lying down-resting |  |  |
| Washing hands and face |  |  |
| Using the toilet |  |  |
| Bathing |  |  |
| Running |  |  |
| Chatting with somebody |  |  |
| Sitting |  |  |
| Counting money |  |  |
| Wearing-taking off shoes |  |  |
| Other: Dying, making peace,<br>doing politics, cutting ribbon |  |  |

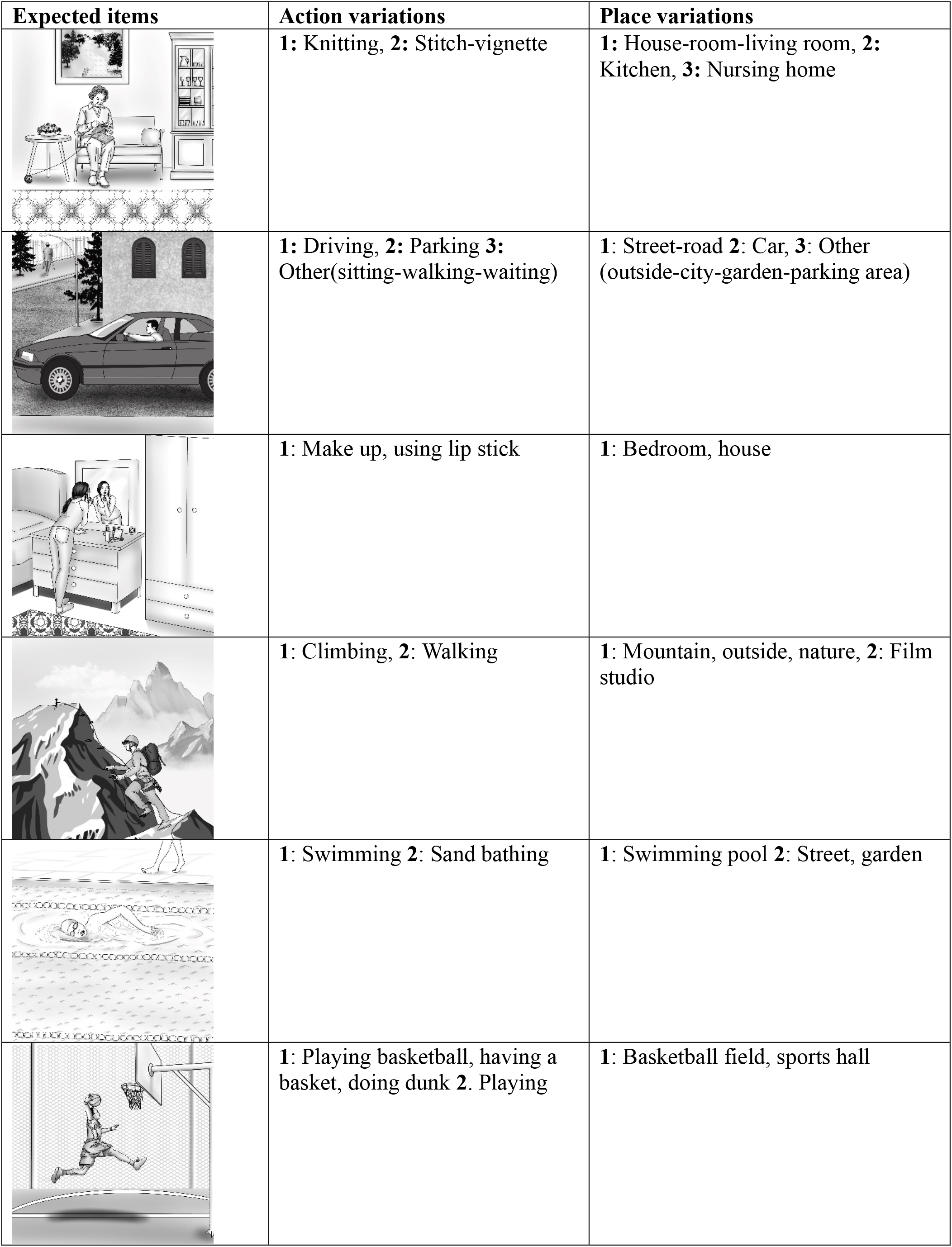
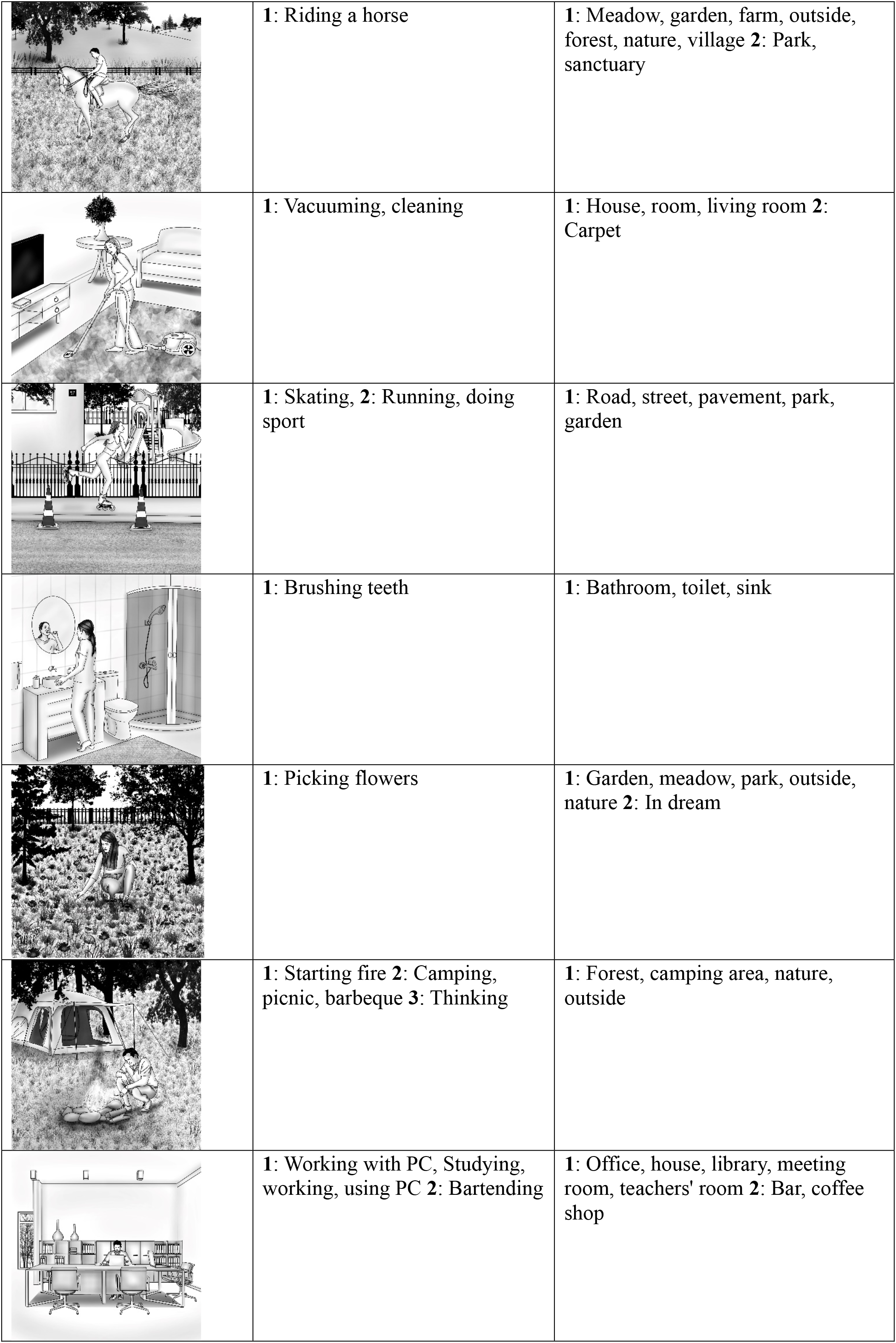
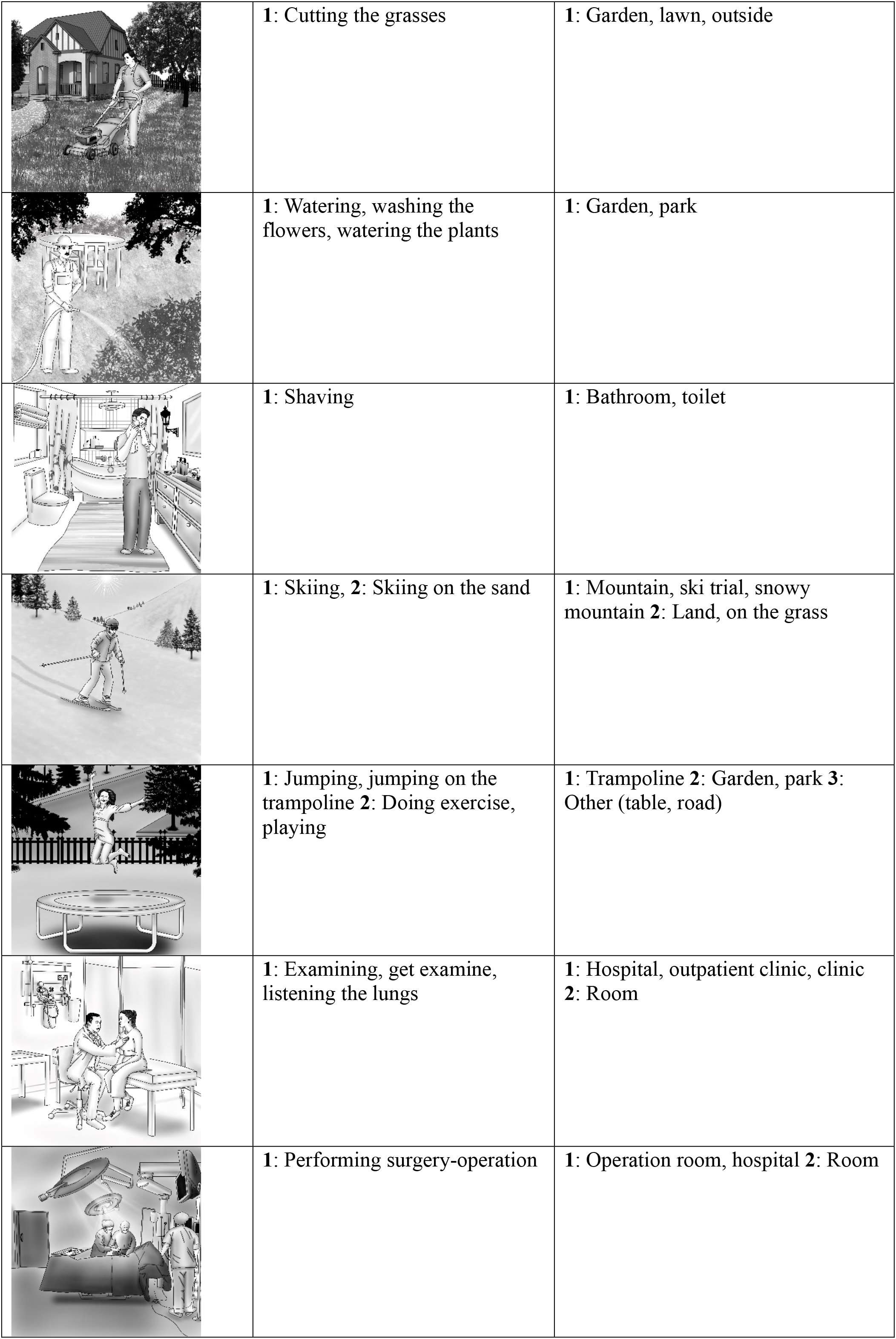
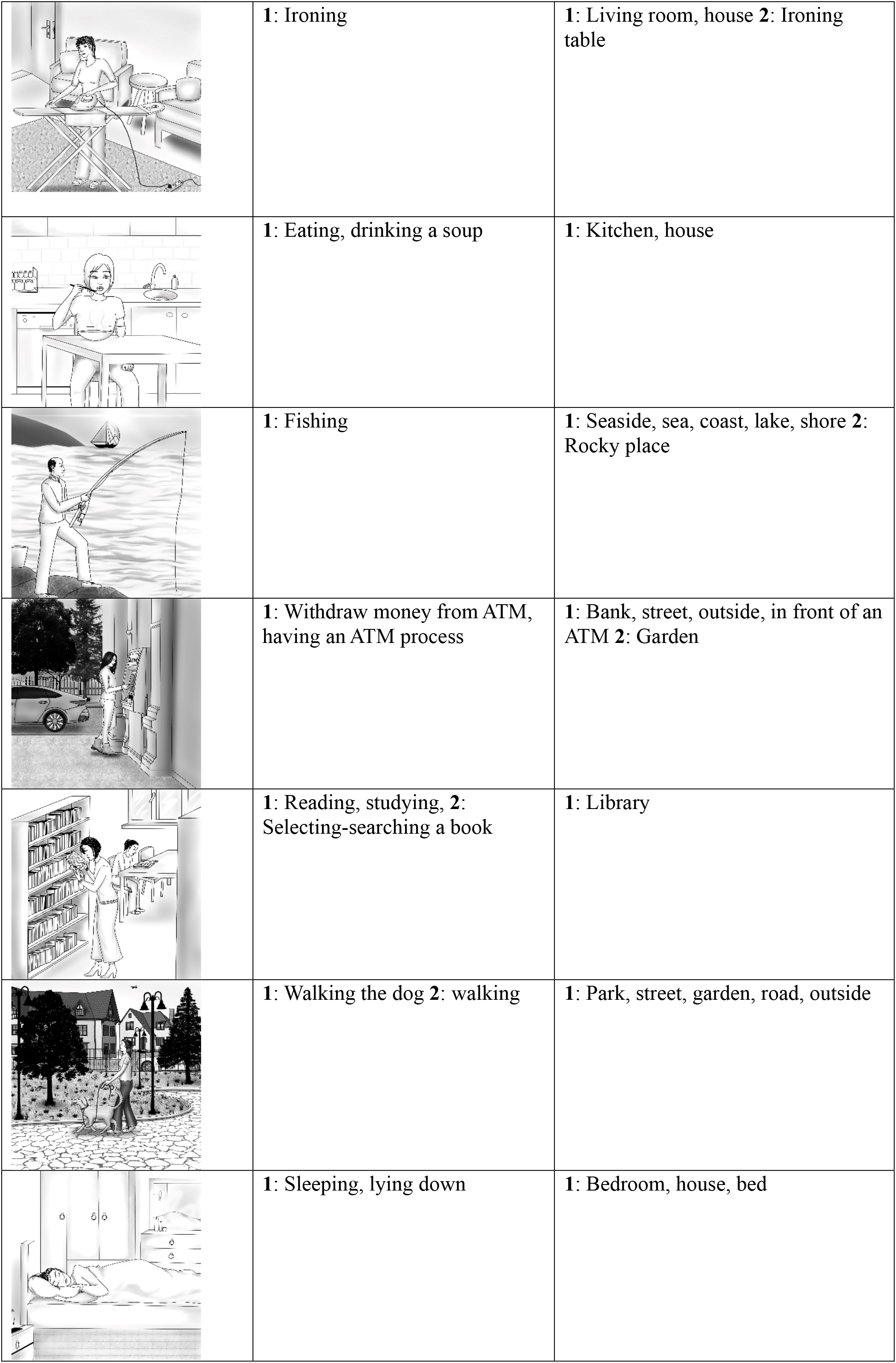
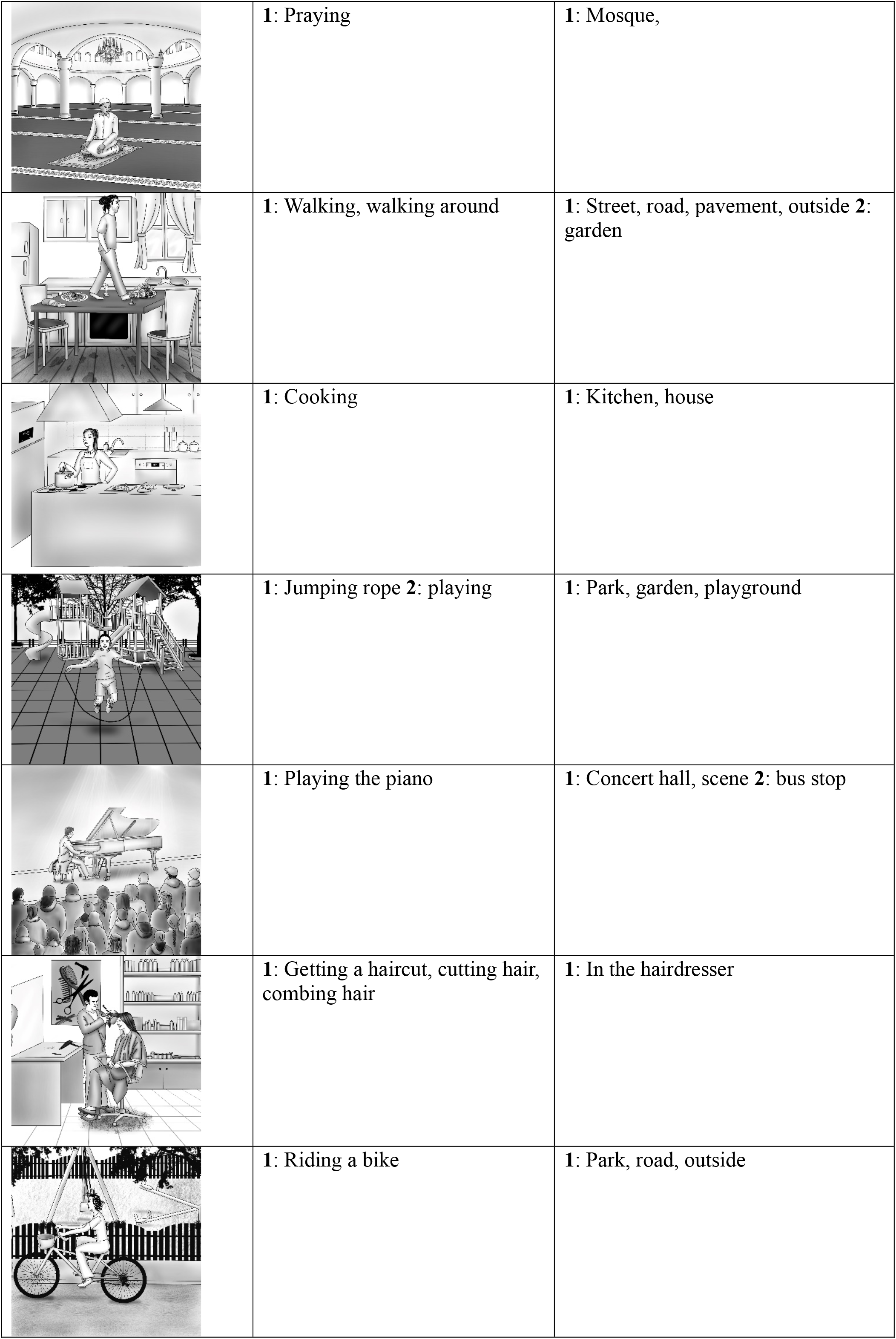
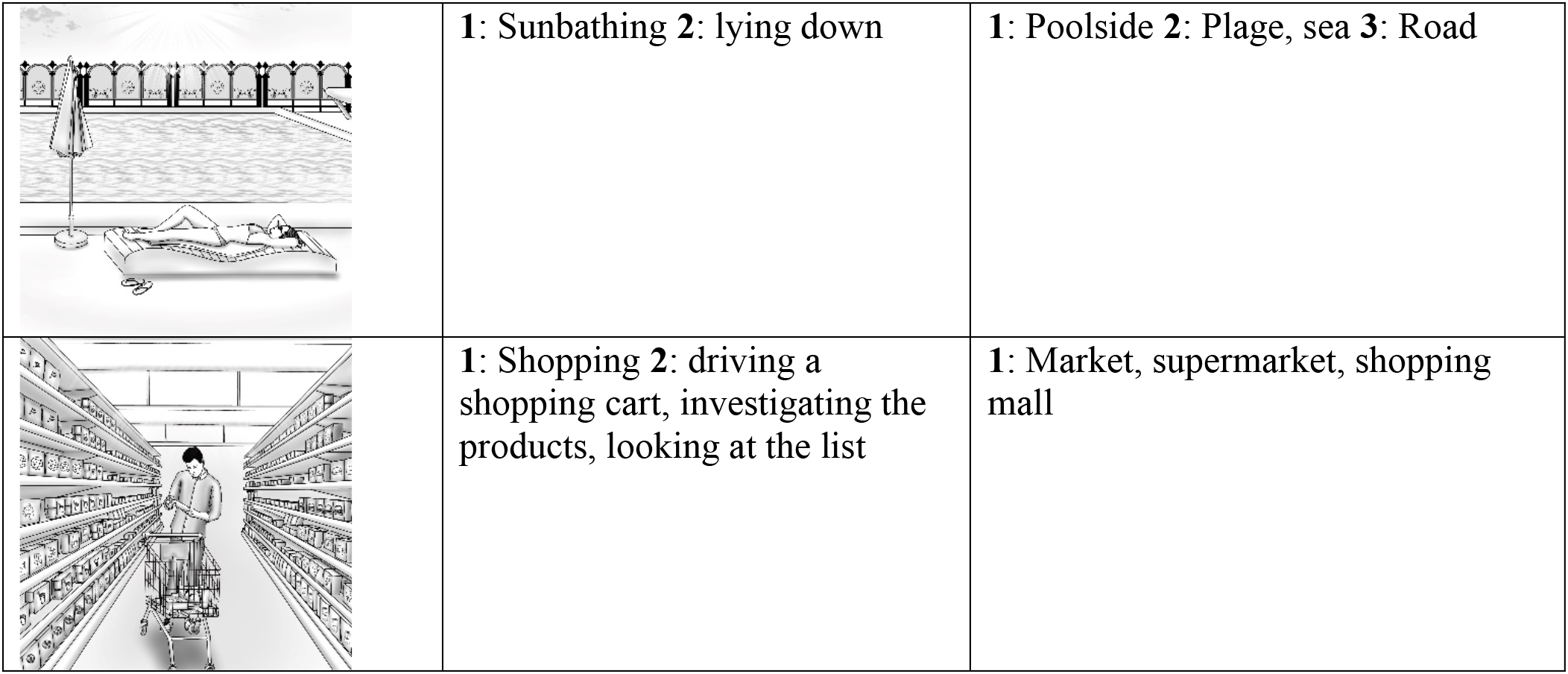

| Expected items | Action variations | Place variations |
| --- | --- | --- |
|  | 1: Knitting, 2: Stitch-vignette | 1: House-room-living room, 2: Kitchen, 3: Nursing home |
|  | 1: Driving, 2: Parking 3: Other(sitting-walking-waiting) | 1: Street-road 2: Car, 3: Other (outside-city-garden-parking area) |
|  | 1: Make up, using lip stick | 1: Bedroom, house |
|  | 1: Climbing, 2: Walking | 1: Mountain, outside, nature, 2: Film studio |
|  | 1: Swimming 2: Sand bathing | 1: Swimming pool 2: Street, garden |
|  | 1: Playing basketball, having a basket, doing dunk 2. Playing | 1: Basketball field, sports hall |
|  | 1: Riding a horse | 1: Meadow, garden, farm, outside, forest, nature, village 2: Park, sanctuary |
|  | 1: Vacuuming, cleaning | 1: House, room, living room 2: Carpet |
|  | 1: Skating, 2: Running, doing sport | 1: Road, street, pavement, park, garden |
|  | 1: Brushing teeth | 1: Bathroom, toilet, sink |
|  | 1: Picking flowers | 1: Garden, meadow, park, outside, nature 2: In dream |
|  | 1: Starting fire 2: Camping, picnic, barbeque 3: Thinking | 1: Forest, camping area, nature, outside |
|  | 1: Working with PC, Studying, working, using PC 2: Bartending | 1: Office, house, library, meeting room, teachers' room 2: Bar, coffee shop |
|  | 1: Cutting the grasses | 1: Garden, lawn, outside |
|  | 1: Watering, washing the flowers, watering the plants | 1: Garden, park |
|  | 1: Shaving | 1: Bathroom, toilet |
|  | 1: Skiing, 2: Skiing on the sand | 1: Mountain, ski trial, snowy mountain 2: Land, on the grass |
|  | 1: Jumping, jumping on the trampoline 2: Doing exercise, playing | 1: Trampoline 2: Garden, park 3: Other (table, road) |
|  | 1: Examining, get examine, listening the lungs | 1: Hospital, outpatient clinic, clinic 2: Room |
|  | 1: Performing surgery-operation | 1: Operation room, hospital 2: Room |
|  | **1: Ironing** | **1: Living room, house 2: Ironing table** |
|  | **1: Eating, drinking a soup** | **1: Kitchen, house** |
|  | **1: Fishing** | **1: Seaside, sea, coast, lake, shore 2: Rocky place** |
|  | **1: Withdraw money from ATM, having an ATM process** | **1: Bank, street, outside, in front of an ATM 2: Garden** |
|  | **1: Reading, studying, 2: Selecting-searching a book** | **1: Library** |
|  | **1: Walking the dog 2: walking** | **1: Park, street, garden, road, outside** |
|  | **1: Sleeping, lying down** | **1: Bedroom, house, bed** |
|  | 1: Praying | 1: Mosque, |
|  | 1: Walking, walking around | 1: Street, road, pavement, outside 2: garden |
|  | 1: Cooking | 1: Kitchen, house |
|  | 1: Jumping rope 2: playing | 1: Park, garden, playground |
|  | 1: Playing the piano | 1: Concert hall, scene 2: bus stop |
|  | 1: Getting a haircut, cutting hair, combing hair | 1: In the hairdresser |
|  | 1: Riding a bike | 1: Park, road, outside |
|  | **1:** Sunbathing **2:** lying down | **1:** Poolside **2:** Plage, sea **3:** Road |
|  | **1:** Shopping **2:** driving a shopping cart, investigating the products, looking at the list | **1:** Market, supermarket, shopping mall |

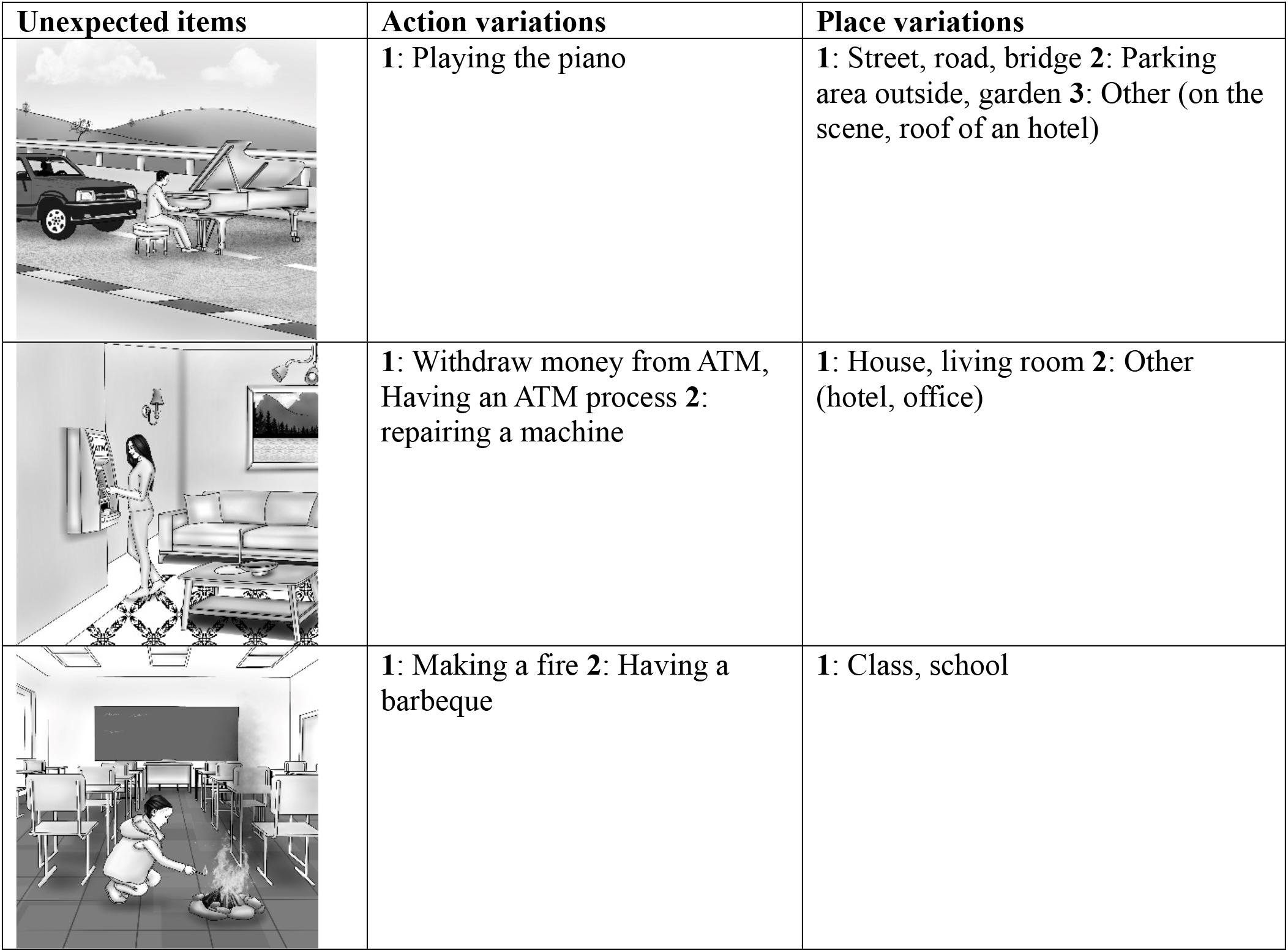
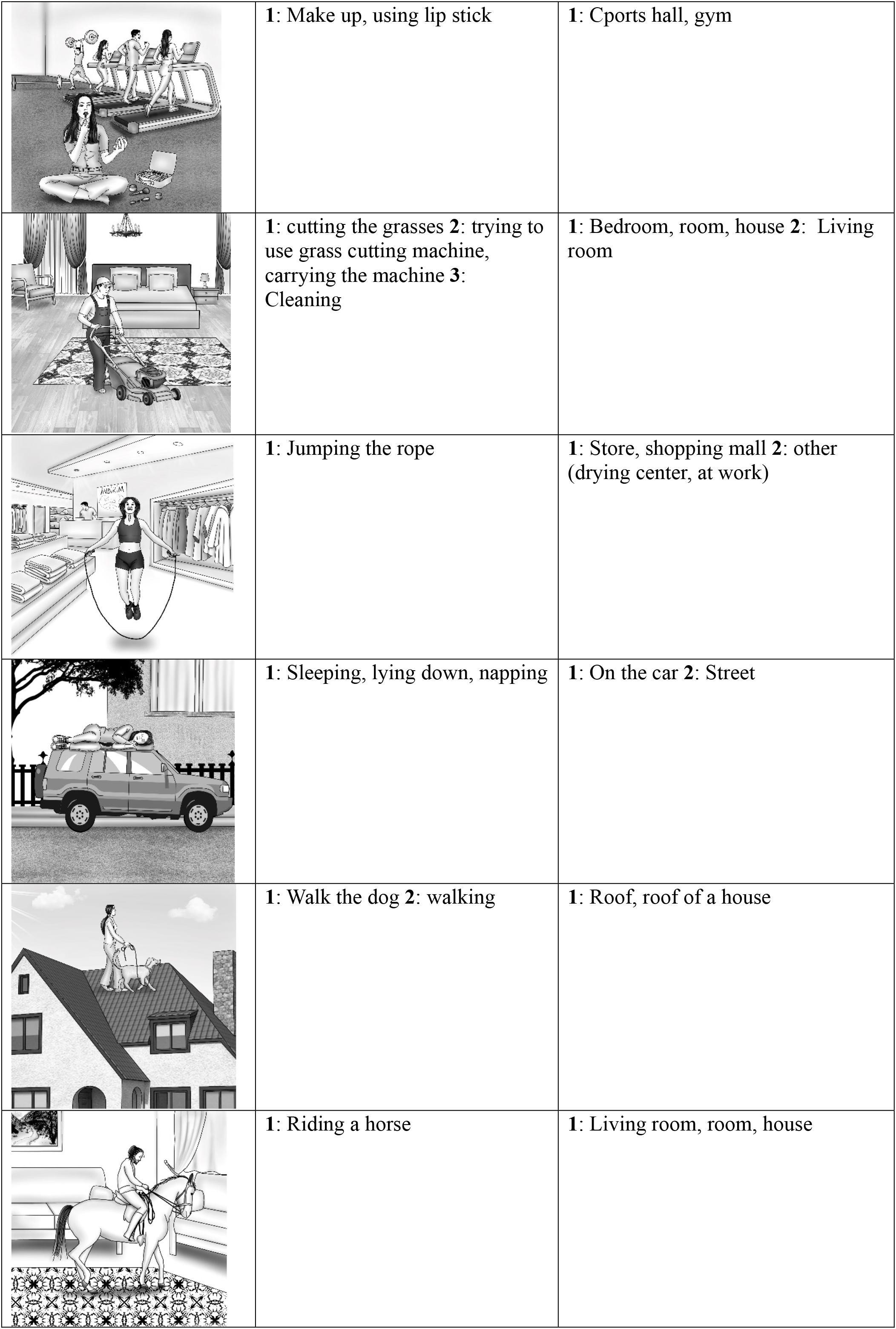
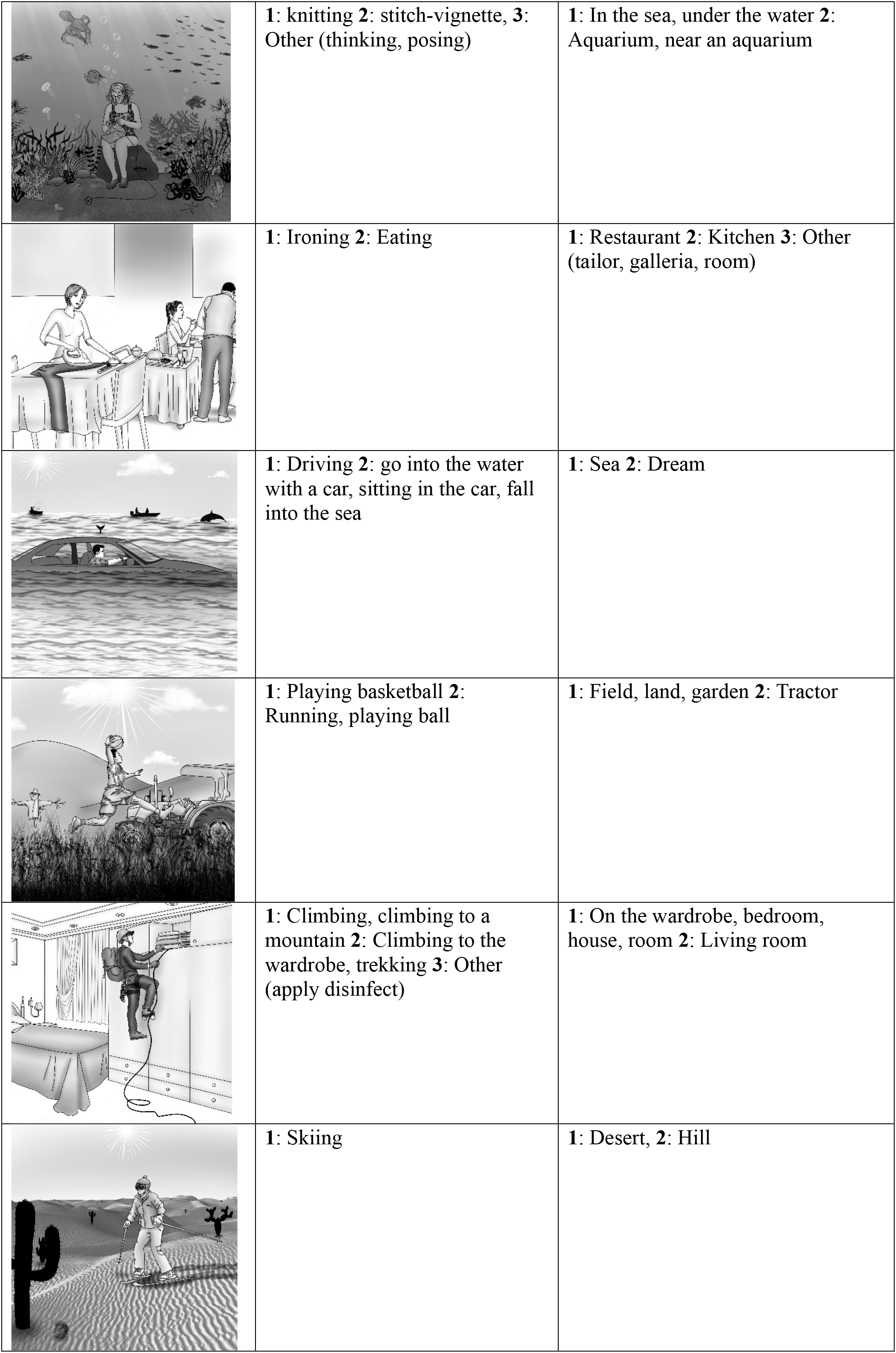
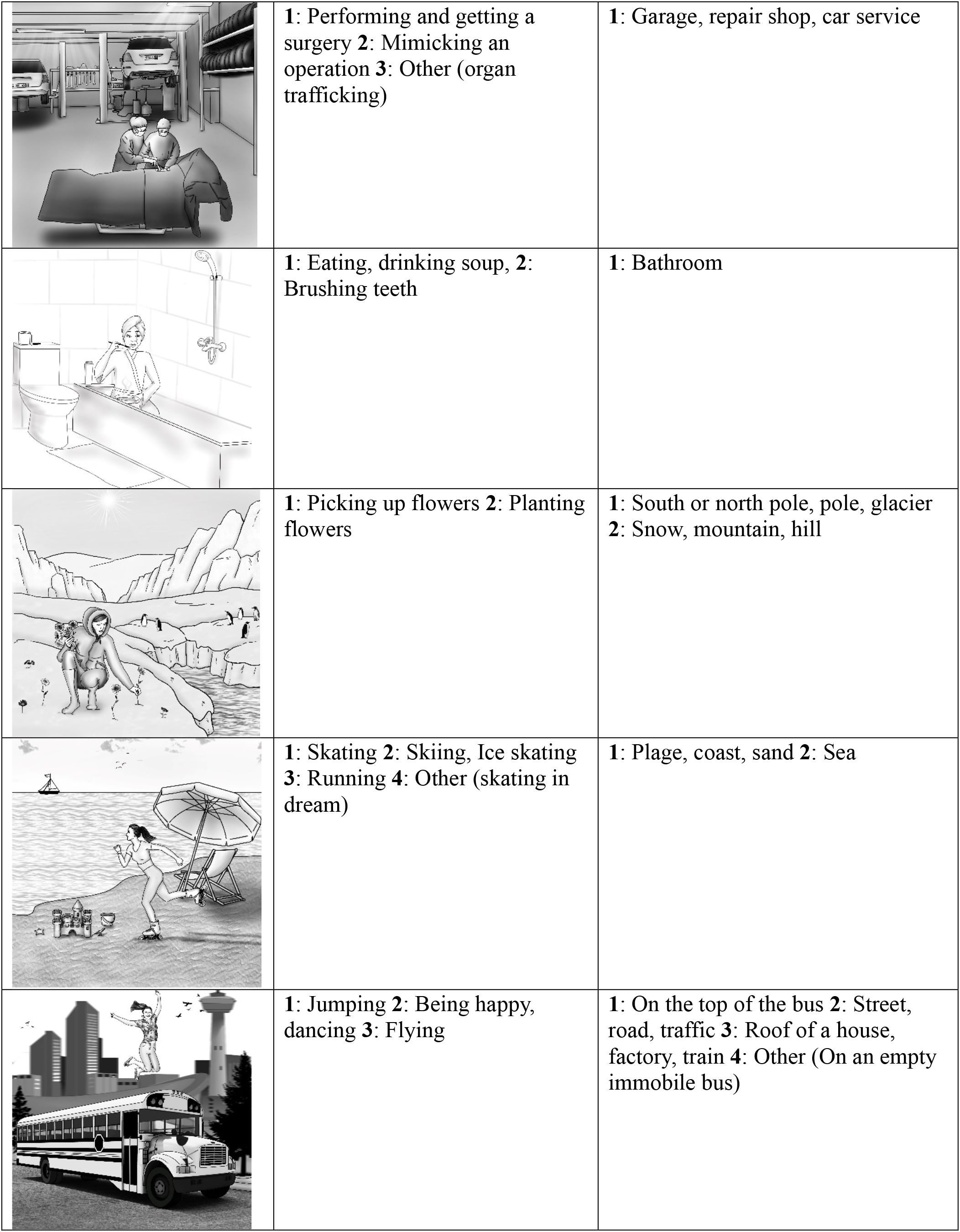
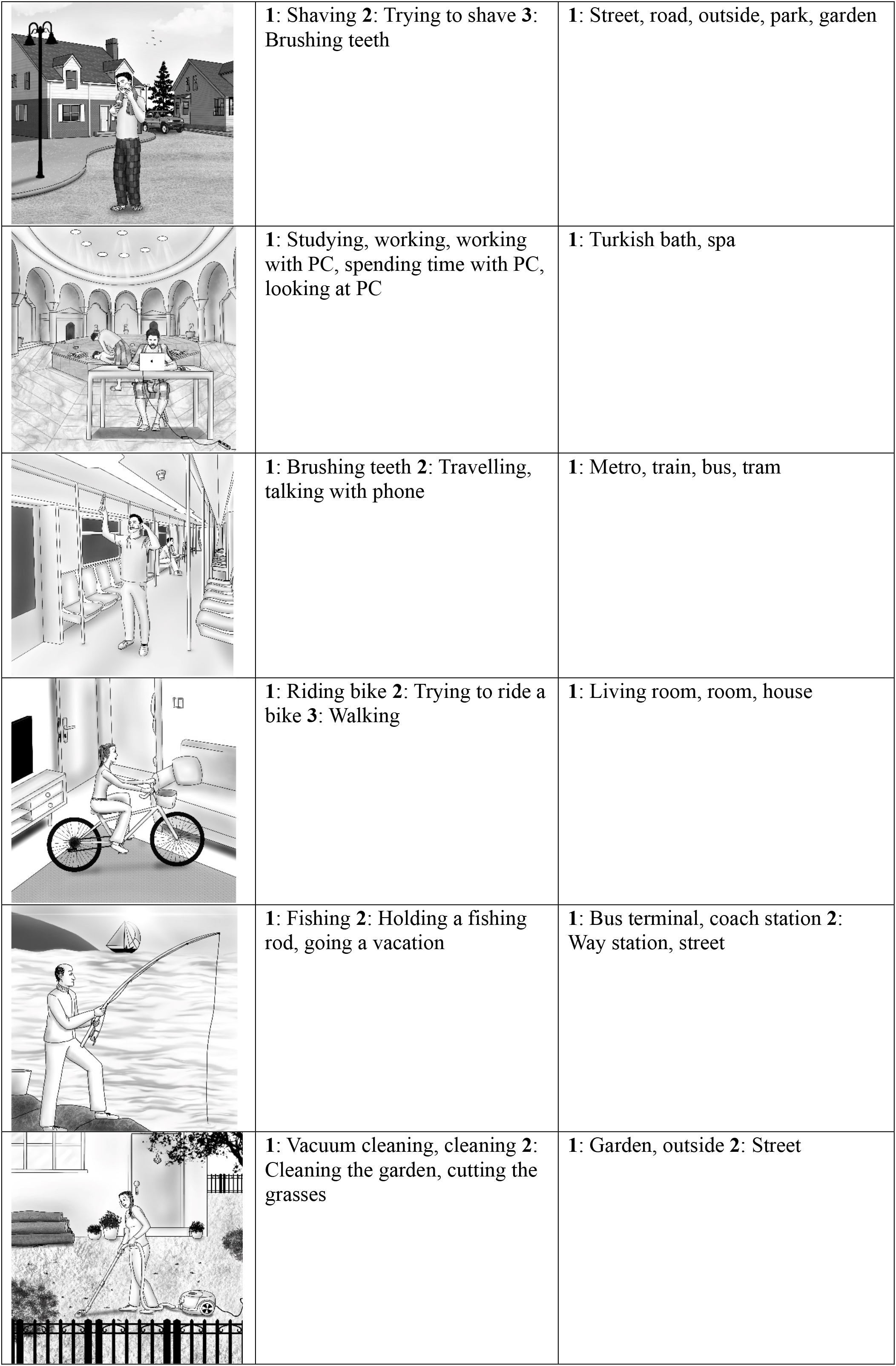
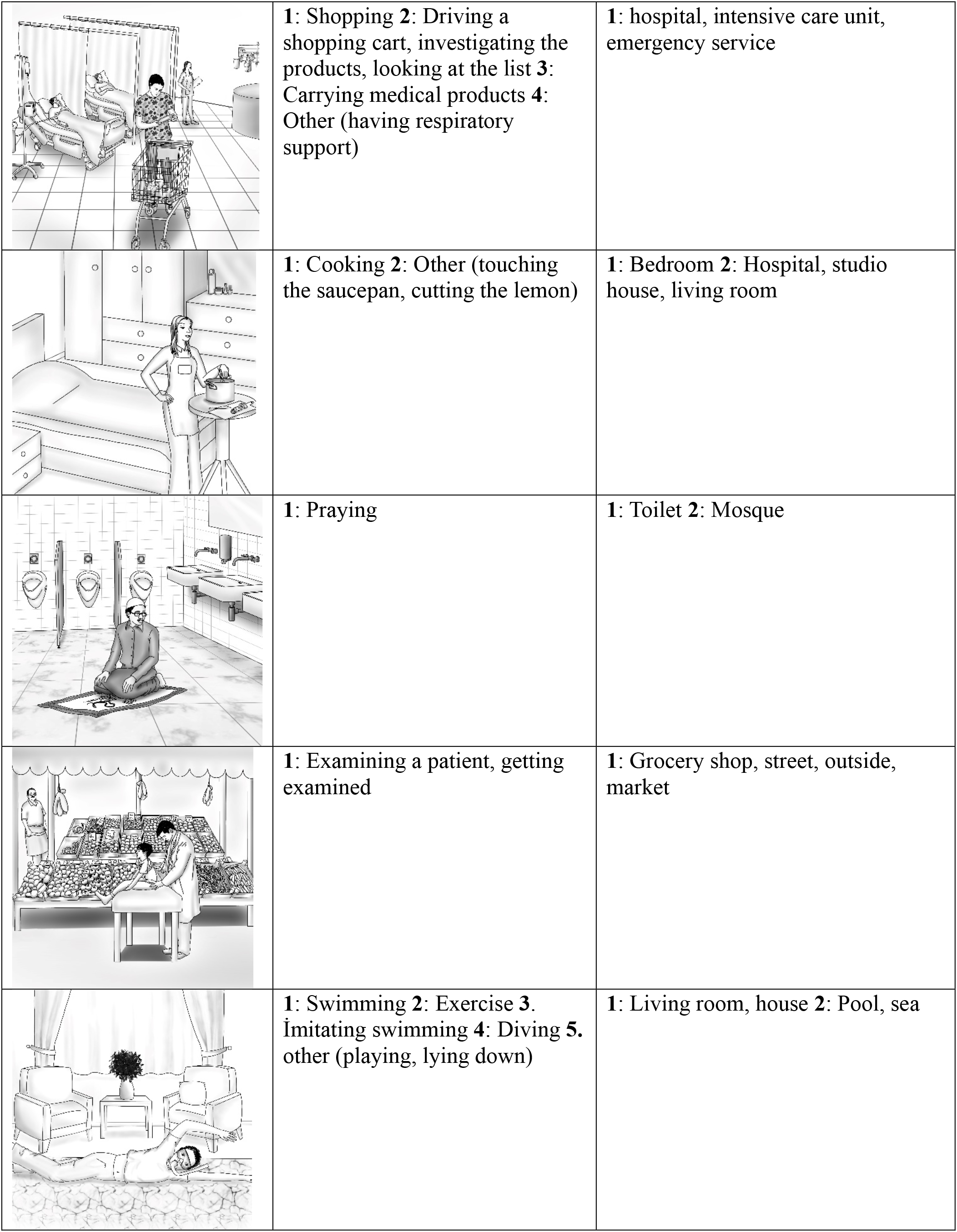
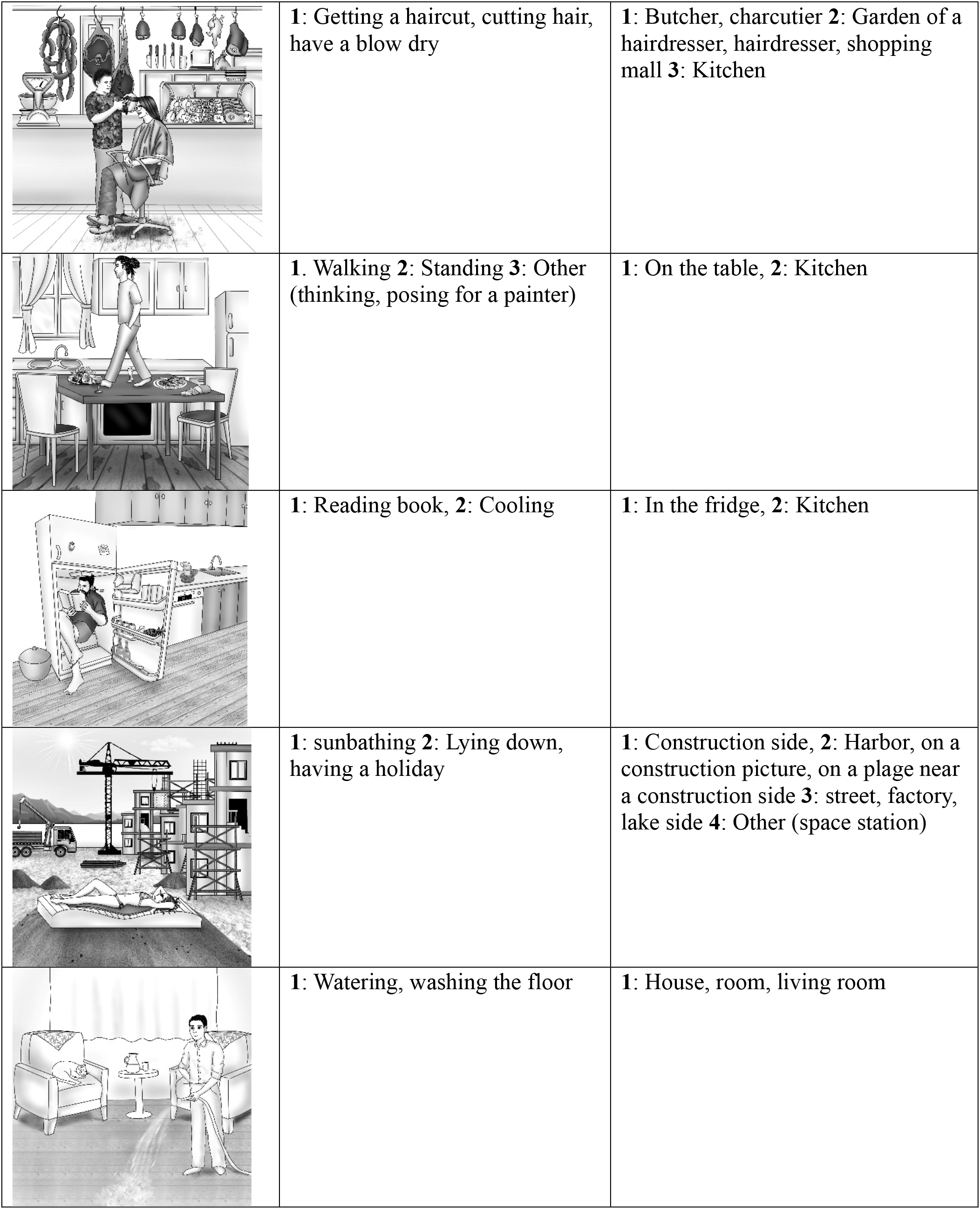

| Unexpected items | Action variations | Place variations |
| --- | --- | --- |
|  | **1:** Playing the piano | **1:** Street, road, bridge **2:** Parking area outside, garden **3:** Other (on the scene, roof of an hotel) |
|  | **1:** Withdraw money from ATM, Having an ATM process **2:** repairing a machine | **1:** House, living room **2:** Other (hotel, office) |
|  | **1:** Making a fire **2:** Having a barbeque | **1:** Class, school |
|  | 1: Make up, using lip stick | 1: Sports hall, gym |
|  | 1: cutting the grasses 2: trying to use grass cutting machine, carrying the machine 3: Cleaning | 1: Bedroom, room, house 2: Living room |
|  | 1: Jumping the rope | 1: Store, shopping mall 2: other (drying center, at work) |
|  | 1: Sleeping, lying down, napping | 1: On the car 2: Street |
|  | 1: Walk the dog 2: walking | 1: Roof, roof of a house |
|  | 1: Riding a horse | 1: Living room, room, house |
|  | **1: knitting 2: stitch-vignette, 3: Other (thinking, posing)** | **1: In the sea, under the water 2: Aquarium, near an aquarium** |
|  | **1: Ironing 2: Eating** | **1: Restaurant 2: Kitchen 3: Other (tailor, galleria, room)** |
|  | **1: Driving 2: go into the water with a car, sitting in the car, fall into the sea** | **1: Sea 2: Dream** |
|  | **1: Playing basketball 2: Running, playing ball** | **1: Field, land, garden 2: Tractor** |
|  | **1: Climbing, climbing to a mountain 2: Climbing to the wardrobe, trekking 3: Other (apply disinfect)** | **1: On the wardrobe, bedroom, house, room 2: Living room** |
|  | **1: Skiing** | **1: Desert, 2: Hill** |
|  | **1:** Performing and getting a surgery **2:** Mimicking an operation **3:** Other (organ trafficking) | **1:** Garage, repair shop, car service |
|  | **1:** Eating, drinking soup, **2:** Brushing teeth | **1:** Bathroom |
|  | **1:** Picking up flowers **2:** Planting flowers | **1:** South or north pole, pole, glacier<br>**2:** Snow, mountain, hill |
|  | **1:** Skating **2:** Skiing, Ice skating<br>**3:** Running **4:** Other (skating in dream) | **1:** Plage, coast, sand **2:** Sea |
|  | **1:** Jumping **2:** Being happy, dancing **3:** Flying | **1:** On the top of the bus **2:** Street, road, traffic **3:** Roof of a house, factory, train **4:** Other (On an empty immobile bus) |
|  | **1: Shaving 2: Trying to shave 3: Brushing teeth** | **1: Street, road, outside, park, garden** |
|  | **1: Studying, working, working with PC, spending time with PC, looking at PC** | **1: Turkish bath, spa** |
|  | **1: Brushing teeth 2: Travelling, talking with phone** | **1: Metro, train, bus, tram** |
|  | **1: Riding bike 2: Trying to ride a bike 3: Walking** | **1: Living room, room, house** |
|  | **1: Fishing 2: Holding a fishing rod, going a vacation** | **1: Bus terminal, coach station 2: Way station, street** |
|  | **1: Vacuum cleaning, cleaning 2: Cleaning the garden, cutting the grasses** | **1: Garden, outside 2: Street** |
|  | **1:** Shopping **2:** Driving a shopping cart, investigating the products, looking at the list **3:** Carrying medical products **4:** Other (having respiratory support) | **1:** hospital, intensive care unit, emergency service |
|  | **1:** Cooking **2:** Other (touching the saucepan, cutting the lemon) | **1:** Bedroom **2:** Hospital, studio house, living room |
|  | **1:** Praying | **1:** Toilet **2:** Mosque |
|  | **1:** Examining a patient, getting examined | **1:** Grocery shop, street, outside, market |
|  | **1:** Swimming **2:** Exercise **3:** Imitating swimming **4:** Diving **5:** other (playing, lying down) | **1:** Living room, house **2:** Pool, sea |
|  | **1:** Getting a haircut, cutting hair, have a blow dry | **1:** Butcher, charcutier **2:** Garden of a hairdresser, hairdresser, shopping mall **3:** Kitchen |
|  | **1.** Walking **2:** Standing **3:** Other (thinking, posing for a painter) | **1:** On the table, **2:** Kitchen |
|  | **1:** Reading book, **2:** Cooling | **1:** In the fridge, **2:** Kitchen |
|  | **1:** sunbathing **2:** Lying down, having a holiday | **1:** Construction side, **2:** Harbor, on a construction picture, on a plage near a construction side **3:** street, factory, lake side **4:** Other (space station) |
|  | **1:** Watering, washing the floor | **1:** House, room, living room |

